# A local inhibitory plasticity rule for control of neuronal firing rate and supralinear dendritic integration

**DOI:** 10.64898/2026.01.20.700499

**Authors:** Daniel Trpevski, Jeanette Hellgren Kotaleski, Matthias Hennig

## Abstract

Inhibitory synapses can control a neuron’s firing rate and also control supralinear dendritic integration. It is not known how inhibitory synapses can learn to perform these functions using only signals available locally at the synaptic site. We study an inhibitory plasticity rule based on the Bienenstock-Cooper-Munro theory in multicompartment models of striatal projection neurons, and show that it can perform these two functions. The rule uses local voltage-gated calcium concentration in the dendrites to regulate inhibitory synaptic strength. We show that, for rate-coded inputs, the rule can achieve precise control of neuronal firing rate after changes in excitatory input rate or excitatory synaptic strength. Additionally, for sparsely-coded inputs that activate localized synaptic clusters in dendrites, the rule can either allow or inhibit the supralinear dendritic response evoked by the clustered excitatory synapses, or equalize the dendritic response arising from different clusters. Finally, we demonstrate the use of learning to inhibit supralinear dendritic integration for solving the nonlinear feature binding problem (NFBP), in tandem with a simple excitatory plasticity rule. We conclude by discussing why the collateral inhibitory synapses between striatal projection neurons could contribute to solving the NFBP with this plasticity rule.

**Author summary:** Neurons are the main cells in the nervous system that process information. They receive signals from the body’s senses—both external and internal—and use them to guide actions such as muscle movement and the regulation of bodily functions. A neuron becomes active when incoming signals excite it strongly enough. But for neurons to work timely, precisely, and reliably, their activity needs to be shaped, modified and controlled. This is done by inhibition, which comes from specialized inhibitory neurons.

In this article we study how inhibition can *learn* to do two of its most basic roles in the nervous system. The first is to help neurons stay responsive across a wide range of input strengths—from very weak to very strong stimulation. For example, neurons in the retina allow vision both in dim starlight and in bright sunlight, even though these conditions differ in brightness by a trillion-fold. Inhibition contributes to handling this huge range by preventing overstimulation of the neurons in bright light. The second role of inhibition is to control strong, local excitations that occur on specific dendritic branches of a neuron. These local excitations can suddenly push a neuron into activity, and inhibition controls whether such excitations are allowed or suppressed.

We use a learning mechanism that is already known to exist for excitatory synapses, but here we apply it to inhibition to explore what it could achieve. The results show that if inhibitory synapses used this same learning rule, they could support the two fundamental roles of inhibition in the nervous system described above.

## Introduction

Inhibitory synapses can significantly control the output of a neuron as well as affect the dendritic computations it performs [1]. For example, inhibition can control a neuron’s responsiveness to excitatory inputs, measured in terms of its output firing rate or spiking probability [2–5]. This responsiveness is described by a neuron’s input-output (IO) function, and inhibition can both shift this function and affect its slope, performing the so-called subtractive and divisive changes to the IO function, respectively. Among other things, such changes to the IO function make the neuron responsive to a much wider range of excitatory input drive (also called expansion of the neuron’s dynamic range). Also, if they are strong enough, inhibitory synapses can suppress supralinear dendritic excitations; alternatively, if they are weak or deactivated, they permit such supralienar excitations in a dendrite [6–9]. Supralinear dendritic excitations are an important computational mechanism for detecting coincident synaptic inputs and are also involved in triggering synaptic plasticity [10]. However, how inhibitory synapses can learn to perform these two types of functions using only local signals available at the synaptic site, such as calcium contentration, is not known.

One of the earliest excitatory synaptic plasticity rules, formulated to explain how orientation selectivity develops in the visual cortex, is the Bienenstock-Cooper-Munro (BCM) rule [11]. It is a phenomenological plasticity rule which suggests that synaptic strength is regulated by the level of activity in the postsynaptic neuron, with activity higher than a threshold level eliciting synaptic strengthening (long-term potentiation, LTP) and activity lower than that threshold resulting in synaptic weakening (long-term depression, LTD). The threshold level is also (slowly) varying and follows the average postsynaptic neuron activity, making the BCM rule the first computational implementation of synaptic metaplasticity. Since its formulation, the key features of the BCM rule have been experimentally verified [12]. These experimental findings include: i) the fact that a single stimulated synapse is bidirectionally modifiable, i.e. can undergo either LTP or LTD depending on the amount of activity, ii) in some synapses the same quantity, such as the intracellular Ca^2+^ influx from NMDA receptors, controls whether LTP or LTD occurs, and iii) some molecular mechanisms responsible for metaplasticity have been proposed, such as activity-dependent regulation of the subunit composition of the NMDA receptors, which changes the future synaptic plasticity outcomes for the same type of stimulation.

Comparatively, the molecular mechanisms underlying inhibitory synaptic plasticity are less well understood, even though many participating molecules have been identified [1, 13, 14]. What is known, however, is that, in addition to homosynaptic plasticity, where stimulating a synapse causes changes in that same synapse, inhibitory plasticity is often heterosynaptic [13]. This means it is evoked by the activity of nearby excitatory synapses, frequently without even activating the inhibitory synapses themselves. Also, in many cases the molecular circuits for inhibitory and excitatory plasticity are similar, sometimes even sharing the same molecules [1, 13, 14] (and, for example, compare the LTP molecular circuitry in [15] and [16]).

Thus, since the molecular machinery exists to implement a BCM rule for excitatory synapses, the same or similar machinery could also exist in some inhibitory synapses. With this rationale, here we study a local inhibitory plasticity rule that is also based on the BCM mechanism, and focus on the functions that can be performed by such inhibitory synapses. We use a multicompartment model of a striatal projection neuron (SPN), and the learning rule uses local calcium activity from voltage-gated calcium channels to modify inhibitory synaptic strengths. We show that such an inhibitory plasticity rule can be used both for regulating a neuron’s output firing rate and for gating supralinear dendritic excitations. Before presenting the results, we mention that a previous version of this inhibitory plasticity rule was first used in [17] for controling supralinear dendritic integration. Here we simplify that rule, investigate it in much greater detail, and interpret and extend its use to additional functions of inhibitory plasticity.

## Results

To show how the inhibitory plasticity rule achieves the two functions, we use two types of simulations. In the first type, the inhibitory plasticity rule is used to regulate a neuron’s firing rate when the neuron is stimulated by rate-coded, distributed excitatory inputs (Fig. 1A). This function can be achieved by using only homosynaptic inhibitory plasticity. In the second type, the rule is used for controlling supralinear dendritic integration evoked by input patterns represented with sparsely-coded, clustered synaptic inputs (Fig. 1B). As will be shown, for this fuction, it is necessary to add heterosynaptic inhibitory plasticity.

**Fig 1.**
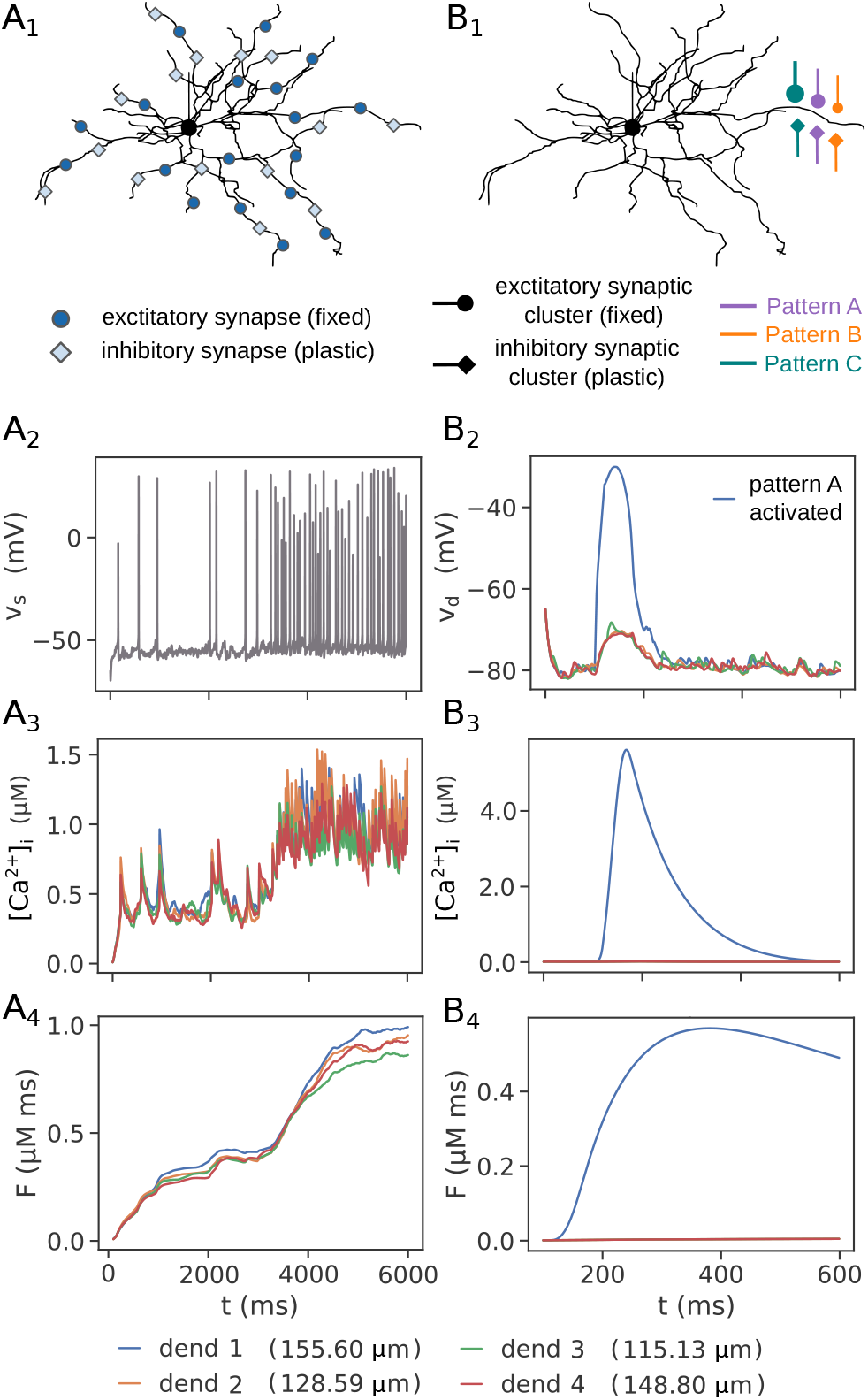
Illustration of the two types of simulations and, for each type, the relationship between voltage, intracellular calcium, and the calcium activity variable *F*. (A_1_) The first type of simulations with rate-coded, distributed synaptic inputs. In these simulations, excitatory synapses are fixed, and inhibitory synapses are plastic. (A_2_ - A_4_) The relation between somatic firing (A_2_) to [Ca^2+^]_i_ (A_3_) and the filtered variable *F* (A_4_) in four different dendrites. At 3 s the exitatory input rate is increased, causing an increase in [Ca^2+^]_i_ and *F*. (B_1_) The second type of simulations with sparsely-coded, clustered synaptic inputs. In most of these simulations excitatory synapses are also fixed, and the inhibitory synapses are plastic. (B_2_ - B_4_) A synaptic cluster of 20 synapses triggers a supralinear dendritic response in one of the dendrites (B_2_), causing an increase in [Ca^2+^]_i_ (B_3_) and *F* (B_4_). Recordings from four different dendrites are plotted for both simulation types.

The inhibitory plasticity rule is based on the BCM rule. In the BCM rule synaptic weights are modified based on some indicator of synaptic activity, and there is a threshold level of this activity below which long-term depression (LTD) is triggered and above which long-term potentiation (LTP) is triggered. It is important that the function determining the weight update changes sign below and above the threshold, and that it becomes 0 for zero synaptic activity [11]. (The function is shown in the respective sections for each type of simulation.) As an indicator of excitatory synaptic activity we use the level of intracellular calcium, [Ca^2+^]_i_, in the dendritic shaft where an inhibitory synapse is located. Since [Ca^2+^]_i_ is a fast signal, we based the plasticity rule on a filtered variable of [Ca^2+^]_i_, called *F*, whose dynamics is slower and “smoother” than that of [Ca^2+^]_i_. It can be thought of as the concentration of a postsynaptic molecule whose activity level is dependent on [Ca^2+^]_i_ and which represents the average level of [Ca^2+^]_i_ in a certain time frame (Eq. 3 in Methods).

An example of the relationship between voltage, [Ca^2+^]_i_ and *F* is given in Fig. 1. Figure 1A_2_-A_4_ shows an example with the setup in Fig. 1A_1_, with distributed, rate-coded inputs. In Fig. 1A_2_, which shows somatic voltage, in the middle of the simulation there is an increase in the excitatory input firing rate, causing an increase in somatic firing. The action potentials propagate in the dendrites, elevating the voltage there and causing increased Ca^2+^ influx (Fig. 1A_3_). The filtered variable *F* also increases as a result (Fig. 1A_4_). Figure 1B_2_-B_4_ shows an example with the setup in Fig. 1B_1_, with clustered, sparsely-coded inputs. In Fig. 1B_2_, the soma does not spike; instead, an excitatory synaptic cluster of 20 synapses positioned in one dendrite is activated, evoking a supralinear, NMDA-dependent dendritic response. This causes increased Ca^2+^ influx localized only in that dendrite (Fig. 1B_3_), and an increase in the variable *F* (Fig. 1B_4_).

In what follows, all simulations of the first type start with a basal level of excitatory input rate or excitatory synaptic strength, after which a change in the inputs is introduced, which affects the output firing rate, as in Fig. 1A. Inhibitory synaptic plasticity then compensates for the change. In the second type of simulations, on the other hand, the soma does not spike, but synaptic clusters in a dendrite are activated, as in Fig. 1B. Inhibitory synaptic plasticity then acts either to i) normalize the responses to all three clusters, or ii) to allow only excitations by some clusters in a dendrite and inhibit the excitations by other clusters in the same dendrite (for example, allow only excitations by pattern A and inhibit those by patterns B and C in Fig. 1B).

### Part I: Control of the neuronal firing rate

#### Dependence of the IO function on inhibitory input rate and inhibitory synaptic strength

We first study what the neuron’s IO function looks like without inhibitory synaptic plasticity. Excitatory and inhibitory synaptic inputs are distributed across the dendrite and are rate-coded, as in Fig. 1A. We consider the influence of two input parameters: i) the frequency of the excitatory inputs, and ii) the synaptic strength of the excitatory inputs. The output firing rate is zero until the excitatory input becomes strong enough, after which the output firing rate linearly depends on both input frequency and synaptic strength (Figs. 2A, B). Inhibitory synaptic plasticity will change the inhibitory synaptic strength, so here we also inspect how the inhibitory synaptic strength affects the IO function: increasing it moves the IO function to the right, i.e. higher excitatory input frequency or synaptic strength is required to reach the firing threshold. This is a so-called subtractive modulation of the input gain, or a shift of the IO function [2]. (The same effect is achieved with an increase in inhibitory input rate, instead of inhibitory synaptic strength, as shown in S1 Fig.)

**Fig 2.**
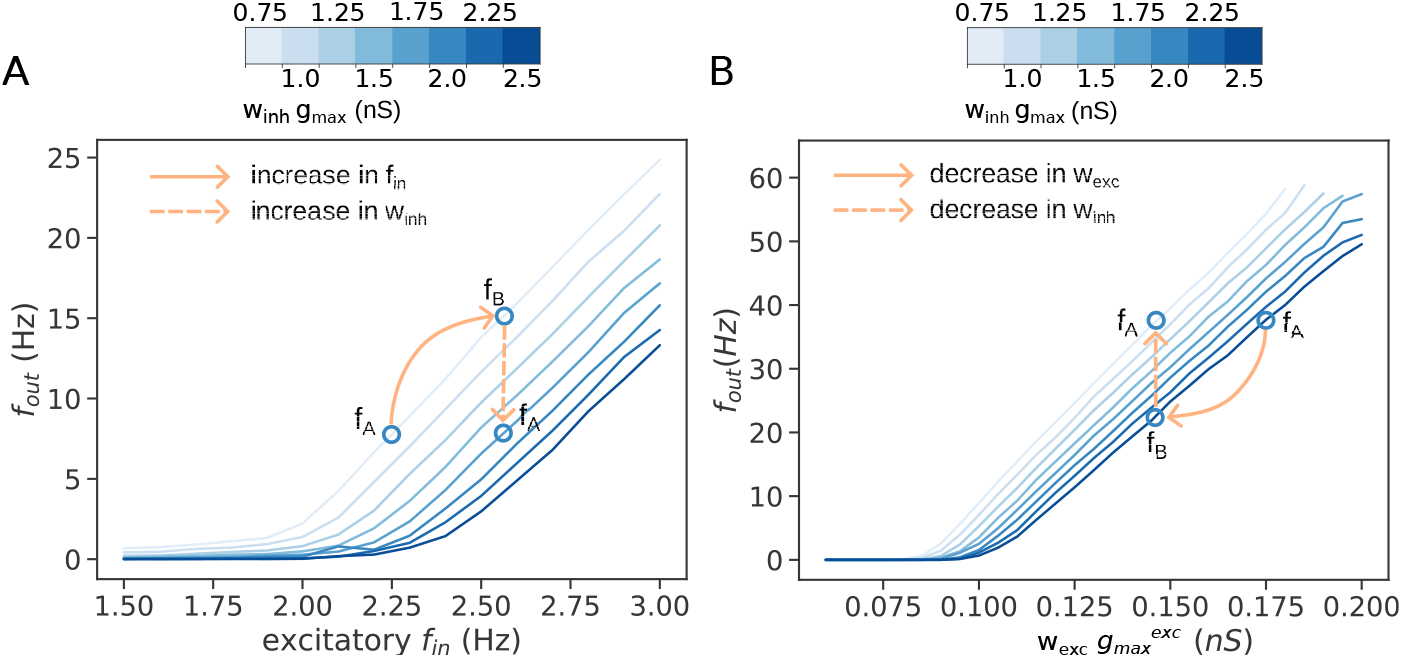
The effect of inhibitory synaptic strength on the IO function. All traces are averages over 50 trials. (A) The IO function showing the neuron’s firing rate versus the input frequency of excitatory inputs. The different lines show the effect of varying the inhibitory synaptic strength. The inhibitory input frequency is 2.3 Hz, and the excitatory synaptic strength is 100 pS. (B) The IO function showing the neuron’s firing rate versus the synaptic strength of excitatory inputs. As in (A), the different lines show the effect of varying the inhibitory synaptic strength. The excitatory and inhibitory input frequency is 2.3 Hz.

Thus, in a scenario where excitatory input rates or excitatory synaptic strengths change (illustrated as solid-line transitions from firing rate *f*_*A*_ to firing rate *f*_*B*_ in Figs. 2A and 2B), inhibitory plasticity should in principle be able to restore the neuronal firing rate to the value before the change in the excitatory inputs (dashed-line transitions from rate *f*_*B*_ to rate *f*_*A*_ in Figs. 2A and 2B).

#### Homosynaptic inhibitory plasticity can restore neuronal firing rate after a perturbation in excitatory input rate or excitatory synaptic strength

In the first type of simulations we use only homosynaptic inhibitory plasticity described with the function (allso called kernel) in Fig. 3, and given with Eqs. 4-6 in Methods. Only the inhibitory synapses are plastic, while excitatory synapses are fixed. Each inhibitory synapse calculates the variable *F* based on the local dendritic [Ca^2+^]_i_ level where it is positioned, and updates its weight according to the plasticity kernel continuously throughout the whole simulation. When *F* is below the threshold *θ*_+_, synapses are weakened, and when *F* is above *θ*_+_, synapses are strengthened. Initially, for each inhibitory synapse the threshold *θ*_+_ is set to the local basal level of *F*. In this way, before any change in the excitatory input, no significant inhibitory plasticity occurs. Once applied, the change in excitatory input rate or synaptic weight affects the value of *F* (as in the example of Fig. 1A_2_-A_4_), causing deviations from *θ*_+_ and non-zero values of the kernel *K*, thus driving inhibitory synaptic plasticity. Additionally, there is a minimal threshold level of *F* for LTD to occur, *θ*_−_ − *c*. Finally, in the first type of simulations, the threshold *θ*_+_ does not change, i.e. there is no metaplasticity. In this way, the threshold *θ*_+_ “encodes” the previous activity level at the location of the inhibitory synapse. It serves as a “set point” which guides the changes in inhibitory synaptic weights in order to restore the same level of neuronal activity before the change in excitatory input.

**Fig 3.**
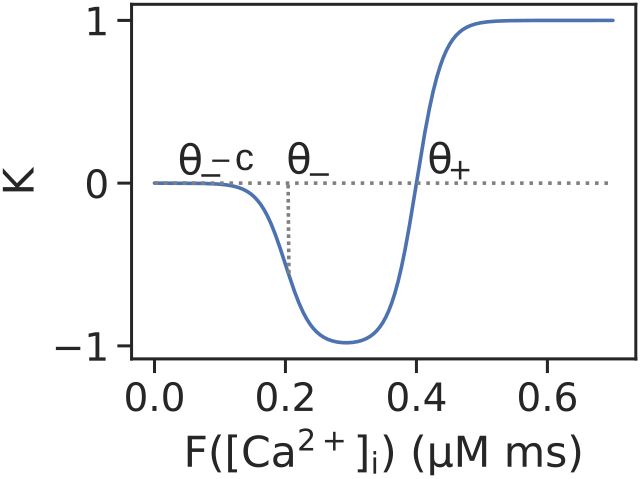
The inhibitory plasticity rule for homosynaptic plasticity used in the first type of simulations (for control of the neuron’s firing rate) and some of the second type of simulations (for equalizing the amplitude of the supralinear dendritic responses). The threshold *θ*_+_ determines the level of *F* below which LTD is triggered and above which LTP is triggered. The threshold *θ*_−_ −*c* is the minimum threshold of *F* for plasticity to occur. (The value *θ*_−_ corresponds to the half-maximum point of the downward slope of the kernel.) The value *c* depends on the steepness parameter in the kernel (see Eq. 6 in Methods). In this kernel the thresholds are fixed, i.e. there is no metaplasticity.

Figure 4 shows how inhibitory plasticity restores the original somatic firing rate after applying an abrupt change in the excitatory inputs. In Figs. 4A_1_-A_3_ the change is an increase of the excitatory input frequency, and in Figs. 4B_1_-B_3_, it is an increase in the excitatory synaptic strength. Both changes can be compensated by inhibitory synaptic plasticity. Figures 4A_1_ and 4B_1_ show the somatic voltage. Figures 4A_2_ and 4B_2_ show the evolution of *F* in four different dendrites, where it can be seen that after the perturbation, inhibitory synaptic plasticity allows the value of *F* to return to the pre-perturbation levels. Figures 4A_3_ and 4B_3_ show the inhibitory weights, some of which have reached a stable level, while some have yet to reach it. Synaptic distance from the soma does not appear to affect the final weight value; however, in Fig. 4B_3_ some proximal inhibitory synapses are notably among the stronger ones, an effect possibly due to the larger proximal amplitude of the backpropagating action potential.

**Fig 4.**
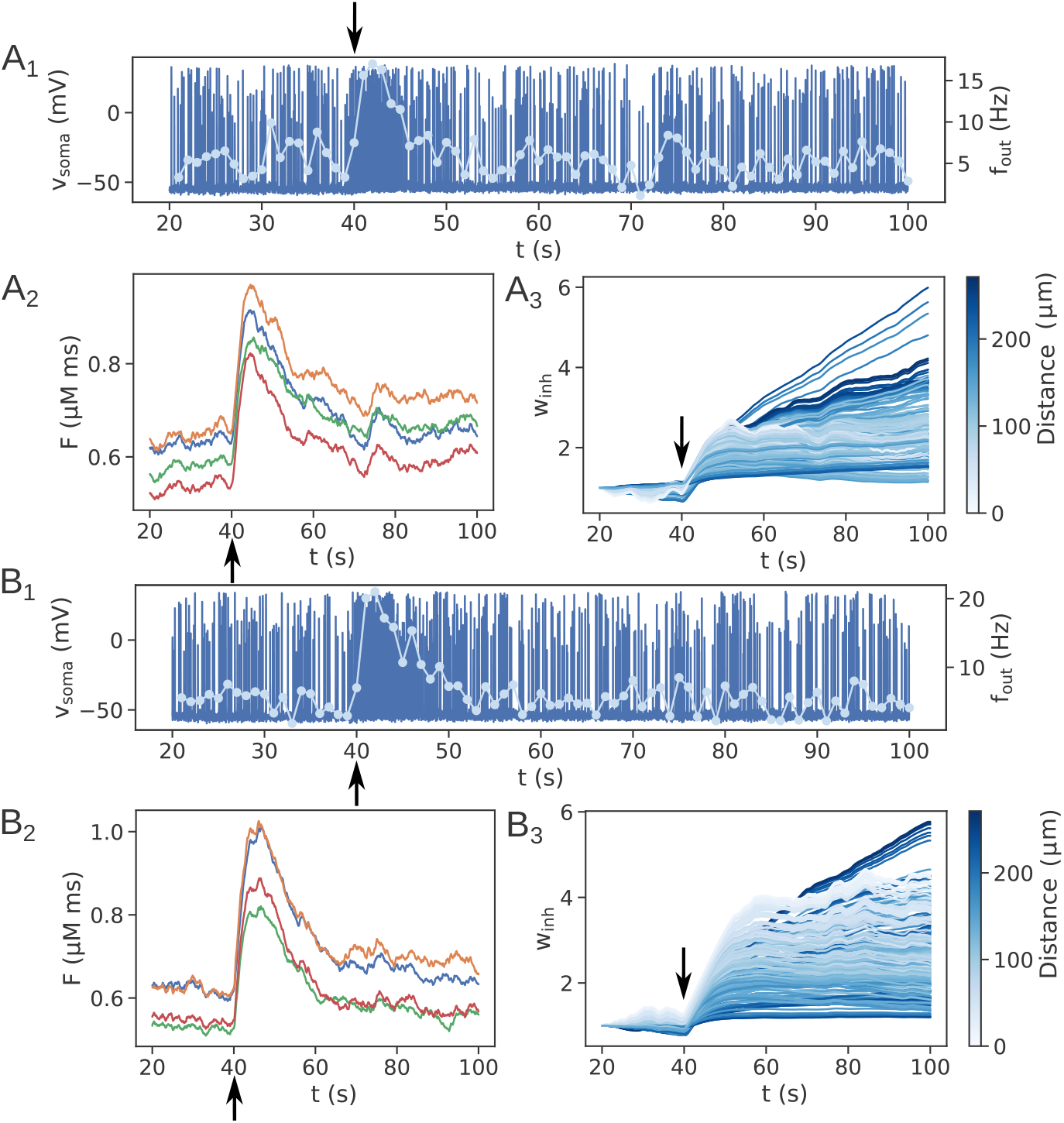
A single run with inhibitory plasticity used to control the neuronal firing rate. (A) A 30% increase in excitatory input frequency is applied. (B) A 25% increase in excitatory synaptic strength is applied. First 20 s where *F* obtains its baseline values are omitted. (A_1_, B_1_) Somatic voltage trace for 100s of simulation time. Change in input is done at 40s (marked with arrows). Markers on the light blue line show the average neuronal firing frequency over the last 1s. (A_2_, B_2_) The filtered [Ca^2+^]_i_ variable *F* recorded from 4 different dendrites at locations of 155 (blue), 128 (orange), 115 (green), and 149 (red) micrometers from the soma. (A_3_, B_3_) The evolution of inhibitory synaptic weights. Line color indicates distance from soma.

We have repeated the simulations in Fig. 4 using different levels of perturbation in the excitatory inputs, averaging over 50 trials for each perturbation level (Fig. 5). The results show that increases in excitatory input frequency and in excitatory synaptic strength can be completely compensated by increasing inhibitory synaptic strength. On the other hand, decreases in input frequency and in excitatory synaptic strength can be compensated by corresponding decreases of inhibitory synaptic strength only up to the point where the decrease in excitatory input still provides enough excitation to drive somatic spiking.

**Fig 5.**
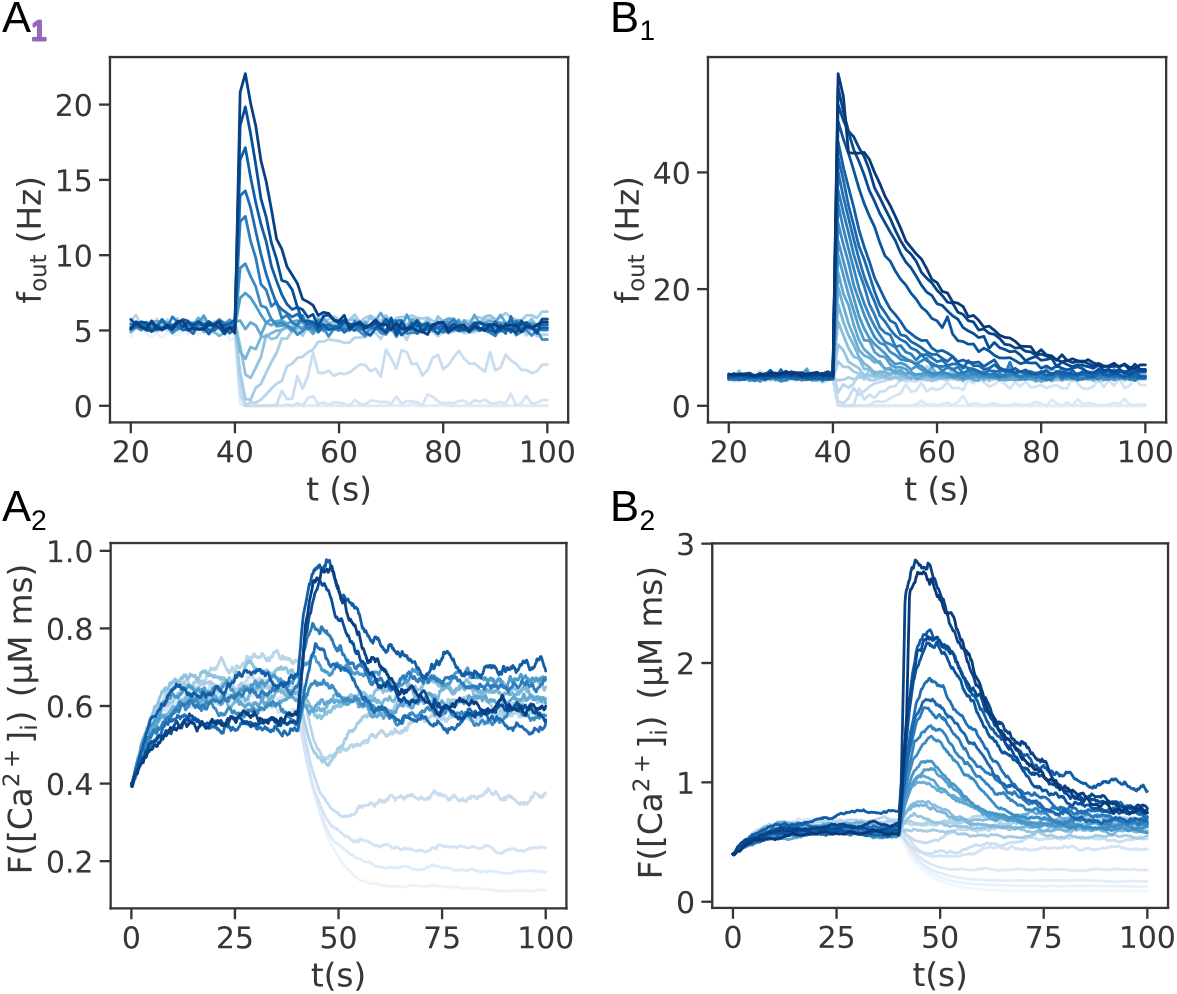
Inhibitory plasticity rule can restore original firing rate for different levels of (A) change in excitatory input frequency and (B) excitatory synaptic strength. Results for each change in input are averages of 50 trials. (A_1_) Excitatiory input frequencies range from 1.5 Hz to 3.0 Hz in steps of 0.1 Hz. Lighter colors are lower input frequencies. Input frequency before the change is 2.3 Hz. Inhibitory input frequency is also 2.3 Hz and remains unchanged. (B_1_) Excitatory synaptic strengths range from 60 pS to 190 pS in steps of 5 pS. Lighter colors are lower synaptic strengths. Synaptic strength before the change is 100 pS. Excitatory and inhibitory input frequency is 2.3 Hz. First 20 s of simulation time where *F* obtains its baseline value are omitted in (A_1_) and (B_1_). (A_2_, B_2_) The variable *F* from a single synapse when varying the excitatory input frequency and excitatory synaptic strength (corresponding to the panels above). Colors indicate the same input values as in (A_1_, B_1_). Traces are from a single trial, and also show the first 20 seconds of simulation time where *F* obtains its baseline value.

In these simulations the thresholds *θ*_+_ and *θ*_−_ are fixed. However, a suitably chosen metaplasticity rule can allow for neuronal firing to settle at a different level compared to the one before the change in excitatory inputs. By changing the activity “set point” encoded with *θ*_+_, a potential mechanism can be provided to neurons to regulate their firing. How neurons determine such set points to control their output firing rate is an open question, so we have not explored such mechanisms further in the first type of simulations [18].

### Part II: Control of supralinear dendritic integration

We also demonstrate that the inhibitory plasticity rule can be used to control supralinear dendritic integration. Specifically, we show this by considering three patterns, A, B, and C, all of which are represented in three dendrites by excitatory synaptic clusters of different sizes (Fig. 6A). In dendrite 1 the strongest pattern is A, represented by a cluster of 20 synapses. In dendrite 2 the strongest pattern is B, and in dendrite 3 the strongest pattern is C. The second strongest pattern in each dendrite is represented by a cluster of 15 synapses, and the weakest pattern is represented by a cluster of 10 synapses. In addition to the excitatory synaptic clusters, each dendrite contains 6 inhibitory synapses, also grouped in clusters, representing each pattern (for a total of 18 inhibitory synapses per dendrite). Patterns are activated sequentially, and are non-overlapping, meaning that when one pattern is activated, both its corresponding excitatory and inhibitory synapses on all dendrites are activated while other synaptic clusters are silent, and synapses representing one pattern are not shared among other patterns. In this setup, the excitatory synaptic weights are fixed, while the inhibitory synaptic weights are plastic. Excitatory synaptic clusters produce NMDA-evoked supralinear dendritic responses of different amplitudes due to the different cluster sizes. Inhibitory synaptic inputs arrive over a longer time window than the excititatory inputs, since they need to suppress long-lasting supralinear dendritic responses.

**Fig 6.**
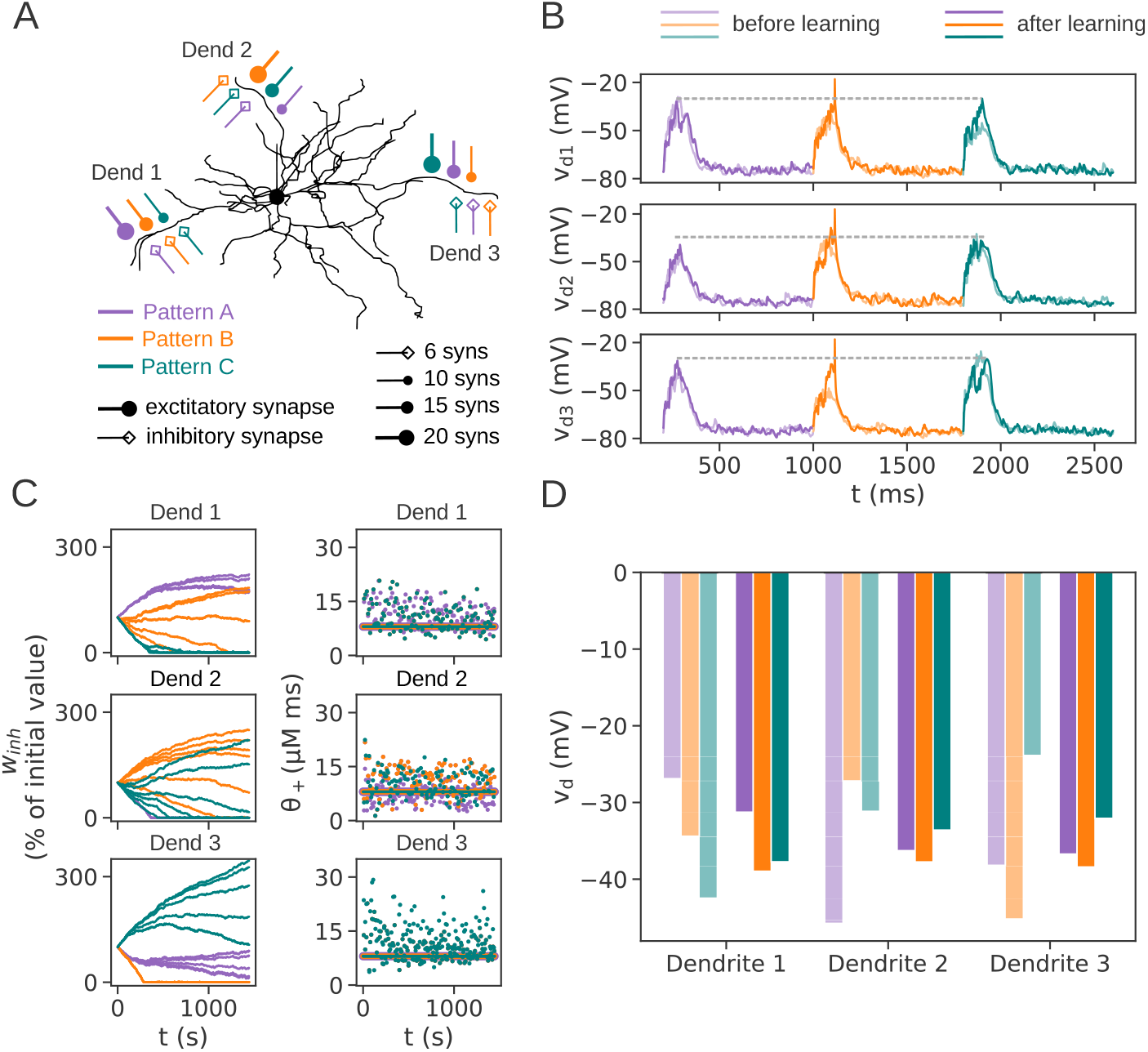
Homosynaptic inhibitory plasticity can equalize the amplitude of supralinear dendritic responses, achieving local gain normalization. (A) The input configuration used in this example. Three dendrites are innervated with excitatory and inhibitory synaptic clusters representing three different patterns, A, B and C. Cluster sizes and identities are indicated in the legend. Excitatory inputs are activated with 3 spikes arriving within a window of 30 ms, while inhibitory synapses are activated with 10 spikes within a window of 100 ms. (B) Dendritic voltage before and after inhibitory synaptic plasticity. In each dendrite, all patterns evoke similar dendritic responses after learning (dashed line provided as a guide for comparison). (C) (Left column) The evolution of inhibitory synaptic weights during learning. In general, the weights for the strongest pattern in each dendrite are increased, those for the weakest pattern are decreased, and those for the middle pattern are either increased or deacreased. (Right column) The maximal value of *F* approaches the fixed threshold *θ*_+_ during learning. *θ*_+_ is shown with lines and max (*F*) with dots. The plots show *θ*_+_ and *F* only for the first synapse in each cluster. Overlapping values of *θ*_+_ are shown with lines with different thickness, while overalpping dots are not shown for clarity. (D) The average dendritic voltage before and after learning. Before learning, the response amplitudes for the three patterns vary significantly, while after learning they are much more similar. Averages are done across 128 trials, with a total of 1800 pattern presentations per trial arriving in random order. The bars show the mean amplitude of the first 10 and the last 10 appearances of a given pattern. Also, in each trial the clusters are placed on three randomly chosen dendrites out of a set of 9 dendrites.

#### Homosynaptic inhibitory plasticity can equalize the amplitude of supralinear dendritic responses

We first examine the effects of homosynaptic plasticity. During homosynaptic plasticity, only the inhibitory synapses which are activated by a pattern are plastic. The other, inactive inhibitory synapses belonging to the other patterns are not plastic for the duration of the activated pattern. (For example, if pattern A arrives, plasticity can be triggered only in the inhibitory synapses representing pattern A, and not in those for patterns B and C.) If we use the same kernel as in the first type of simulations, shown in Fig. 3, the result is analogous to controling the neuronal firing rate: inhibitory synapses are adjusted to achieve the same local voltage evoked by the three different clusters (Fig. 6). This can be seen in Fig. 6B, where after learning the voltage elevations in each dendrite evoked by the three different patterns are very similar, while before learning they are different (due to the different sizes of the excitatory clusters). Each pattern that produced voltage elevations higher than the “set point” determined by the threshold *θ*_+_ had its inhibitory synapses strengthened, while the patterns producing lower voltage elevations had their inhibitory synapses weakened (Fig. 6C, left). Also, analogously to Figs. 5A_2_, B_2_, as learning progressed, the evoked value of the filtered variable *F* showed a trend of moving closer towards the threshold (set point) *θ*_+_, which is kept fixed without metaplasticity (Fig. 6C, right). The results are also confirmed by the average dendritic response to each pattern before and after learning, obtained from 128 trials (Fig. 6D). Before learning, the dendritic responses to each pattern are different due to the different sizes of the excitatory clusters that evoke them. After learning, the dendritic responses to the three patterns in a dendrite become very similar in amplitude. This type of local gain normalization at the level of a dendrite could be useful to make the dendrite and the neuron responsive to which features are active, irrespectively of the size of the synaptic clusters that represent them.

To summarize, both the local, dendritic gain normalization shown in this section and the global, somatic gain normalization in the previous section are achieved with the BCM kernel in Fig. 3 because this kernel makes inhibitory plasticity act as a negative feedback loop in the system: inhibition adapts to match the levels of excitation in order to achieve a “set point” of activity.

#### Homosynaptic inhibitory plasticity can gate or inhibit supralinear dendritic responses, but shows instability in the synaptic weights

Homosynaptic inhibitory synaptic plasticity can also be used to achieve a different function - to permit or inhibit supralinear dendritic responses. To achieve this, we use an inverted BCM kernel, shown in Fig. 7A. This kernel turns inhibitory plasticity into a positive feedback loop onto dendritic voltage and the variable *F*, so to achieve stability of the inhibitory weights during learning, it is necessary to implement metaplasticity. Metaplasticity modifies the value of *θ*_+_, which is also called the sliding threshold in the BCM rule. There are different ways of implementing a sliding threshold, and we have chosen a simple way where *θ*_+_ follows the maximal value of *F* achieved during each pattern presentation (Fig. 7B, Eqs. 8 and 9 in Methods). When *θ*_+_ reaches that value (usually after many update steps), synaptic plasticity stops, because the kernel is 0 at the point of *θ*_+_, making no changes to the synaptic weights.

**Fig 7.**
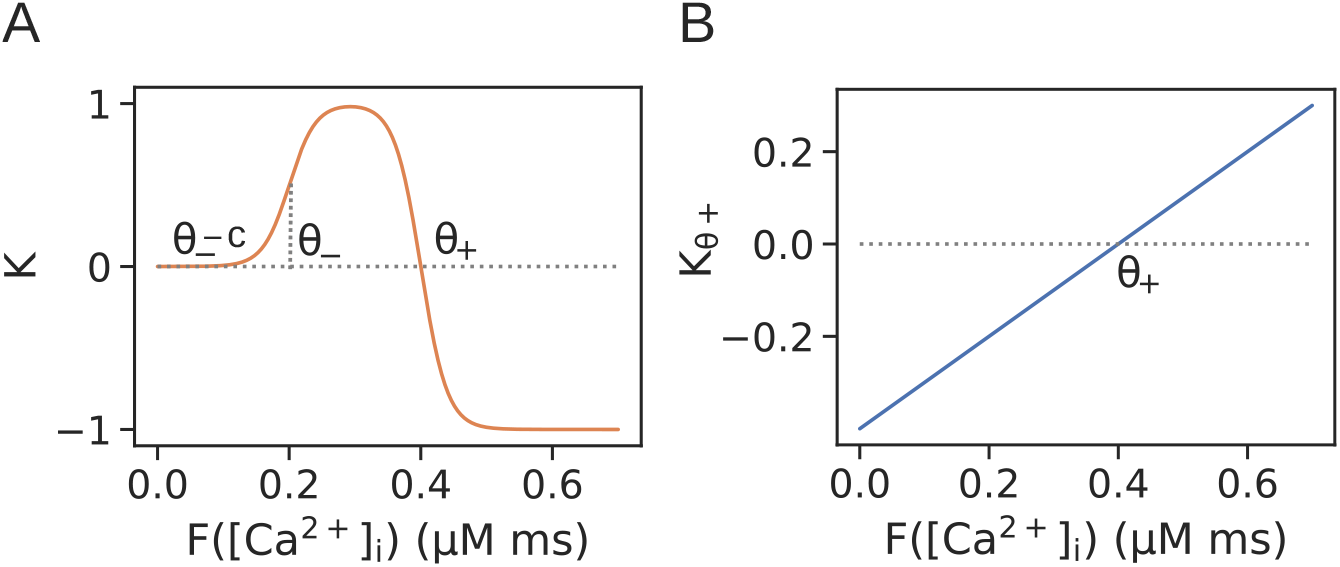
The inhibitory plasticity rule for homosynaptic plasticity used in some of the second type of simulations (for gating or inhibiting supralinear dendritic integration). (A) The plasticity kernel is inverted compared to the kernel in Fig. 3, meaning that for levels of *F* below the threshold *θ*_+_, LTP is triggered, and for levels above *θ*_+_, LTD is triggered. *θ*_−_ − *c* is the minimum level of *F* necessary for plasticity to occur, as in Fig. 3. In this kernel, the threshold *θ*_+_ changes to follow the most recent levels of the calcium activity *F* at the synapse’s location according to the metaplasticity kernel in (B). (B) The metaplasticity kernel describing how *θ*_+_ is updated for the inverted kenrel in (A) (The metaplasticity kenrel is given with Eq. 9 in Methods). Note that as *θ*_+_ changes, the intersection with the x-axis changes.

The results of homosynaptic inhibitory plasticity using the inverted BCM kernel are shown in Fig. 8. Figure 8B shows the voltage elevations in the three dendrites evoked by each of the three patterns, before and after inhibitory synaptic plasticity. Before plasticity, each dendrite responds supralinearly to each pattern, with an amplitude proportional to the number of synapses in the pattern’s cluster. After plasticity, the two strongest patterns in dendrites 1 and 2 are disinhibited, while the remaining pattern is slightly more inhibited. In dendrite 3, only the strongest pattern dendrite 3 is disinhibited, while the other two patterns are inhibited. This can also be seen from the evolution of the inhibitory synaptic weights in Fig. 8C: the synapses for the two strongest patterns in dendrites 1 and 2 are weakened, while those for the weakest pattern are strengthened. Similarly, in denrite 3 only the synapses for the strongest pattern are weakened, and those for the other two patterns are strenthened. The inhibitory weights stop increasing further when the threshold *θ*_+_ becomes equal to the maximal value of *F* (Fig. 8C, right column). This particular simulation run, however, is not representative of the most common outcome with the parameters settings used in Part II of the results. The final outcome of which patterns are disinhibited and which ones are inhibited depends on the initial value of the threshold *θ*_+_. When repeating the learning simulations with homosynaptic inhibitory plasticity for 128 trials, on average the two strongest patterns are disinhibited, while the weakest pattern is inhibited. This can be seen in the average change in dendritic depolarization, *δ*, obtained by averaging across all trials (Fig. 8D).

**Fig 8.**
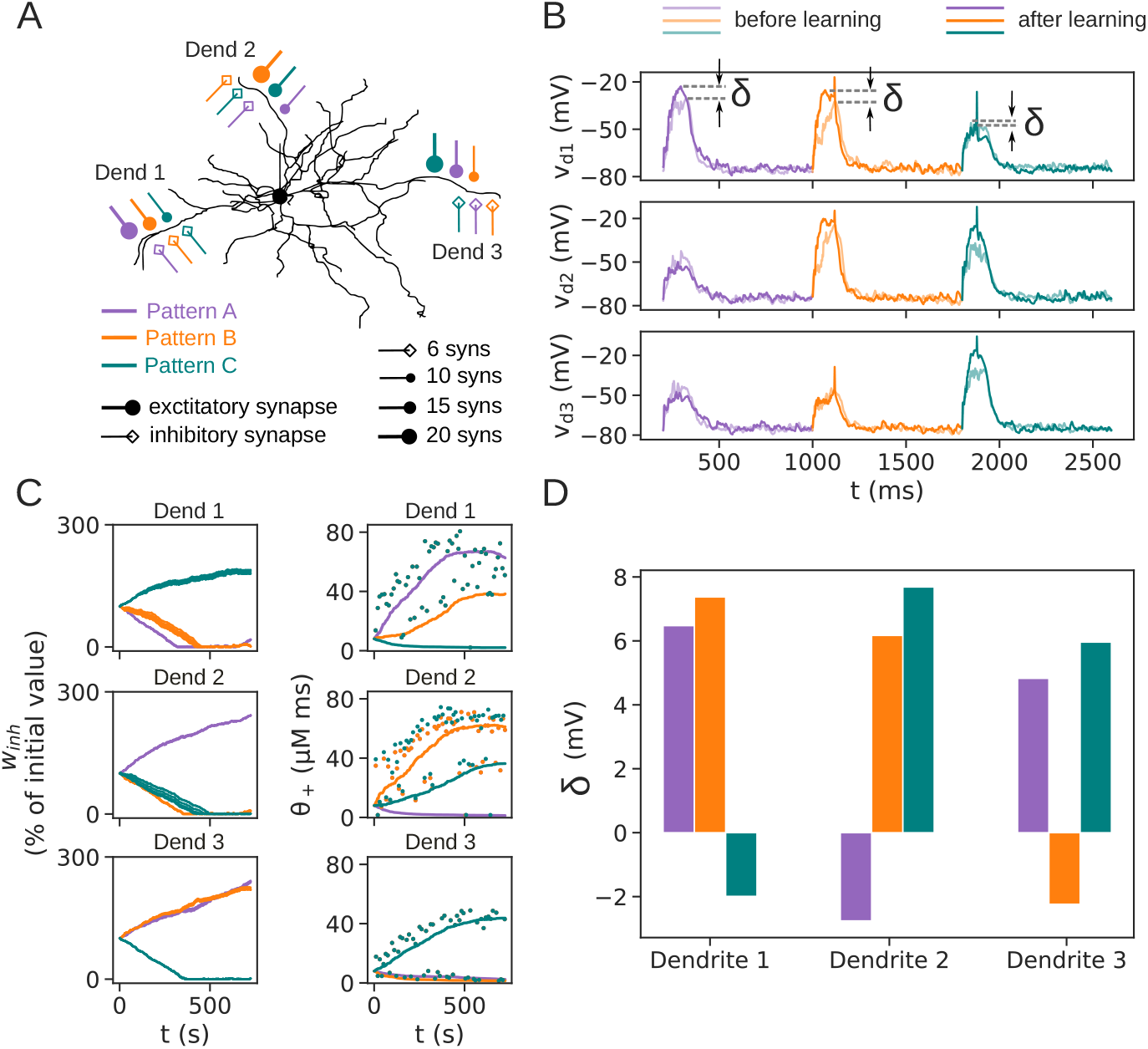
Homosynaptic inhibitory plasticity can gate or inhibit supralinear dendritic responses. (A) The input configuration is the same as in Fig. 6. (B) Voltage traces for each pattern in all three dendrites before and after learning. (C) (Left column) The evolution of inhibitory synaptic weights during learning. The synapses corresponding to the strongest pattern in dendrite 3, and the two strongest patterns in dendrites 1 and 2 are weakened, while the synapses for the remaining patterns are strengthened. (Right column) The evolution of the threshold *θ*_+_ and the maximal value of *F* during learning. *θ*_+_ is shown with lines and max (*F*) with dots, only for the first synapse in each cluster (ovelapping dots are not highighted). When *θ*_+_ reaches the corresponding level of *F*, inhibitory synapses are no longer strengthened. (D) The average change in dendritic depolarization after learning. Positive values indicate increased dendritic responses, and negative values indicate reduced dendritic responses. Results are averages of 128 trials, where in each trial the clusters are placed on three randomly chosen dendrites out of a set of 9 dendrites. Also, in each trial there are a total of 900 pattern presentations arriving in random order. The difference is calculated between the mean of the last 10 and the first 10 appearances of a given pattern.

We briefly note here that in order for patterns to be disinhibited, it is necesarry that the threshold *θ*_+_ is initially not set very high above the evoked values of *F*. If it were set too high, all synaptic activity will be situated below the threshold (the region between *θ*_+_ and *θ*_−_ in Fig. 7A, where the inverted kernel is positive), causing further strengthening of the inhibitory synapses, and inhibiting all patterns. Instead, the threshold *θ*_+_ should be set to a value within the range of the initially evoked values of *F*.

Finally, we highlight that the simulations in Fig. 8 have been run for half the amount of time compared to the similar simulations in Figs. 6, 10 and 11 (only 900 patterns are used versus the usually used 1800 patterns). This is because homosynaptic plasticity with the inverted BCM kernel in Fig. 7A exhibits instability of the synaptic weights for longer simulation times (S2 Fig). This happens because of the positive feedback loop that inhibiton adds to the system, the effects of which are already slightly visible in the top two panels of Fig. 8C, and because *θ*_+_ is just a single point where the kernel has the value 0. As the strongest pattern in dendrite 1 is being disinhibited during learning, the thresholds *θ*_+_ of the inhibitory synapses move towards the increasingly higher maximal values of *F* (purple line going upwards in Fig. 8C, top right panel). If *θ*_+_ were to become equal to *F*, learning would stop completely and the synaptic weights would be stable. However, since *θ*_+_ is just a single point, equality between *θ*_+_ and *F* is difficult to achieve in practice because any small variation in *F* can easily place it slightly below *θ*_+_ (as can be seen with the green dots falling below the purple line in Fig. 8C after 500s, top right panel). Once this happens, the inhibitory weights will be strengthened (Fig. 8C, purple synapses in top left panel at the very end of the simulation), initiating the positive feedback loop of reducing the supralinear dendritic response and the evoked value of *F*, thus keeping it in the positive range of the kernel between *θ*_+_ and *θ*_−_ −*c* where weights are increased even further. This will continue to happen for all future presentations of the strongest pattern, eventually inhibiting it as well. For long simulation times this happens in all three dendrites (S2 Fig, panels C and D). As will be seen in the next section, heterosynaptic plasticity is one way to remove this instability.

**Fig 9.**
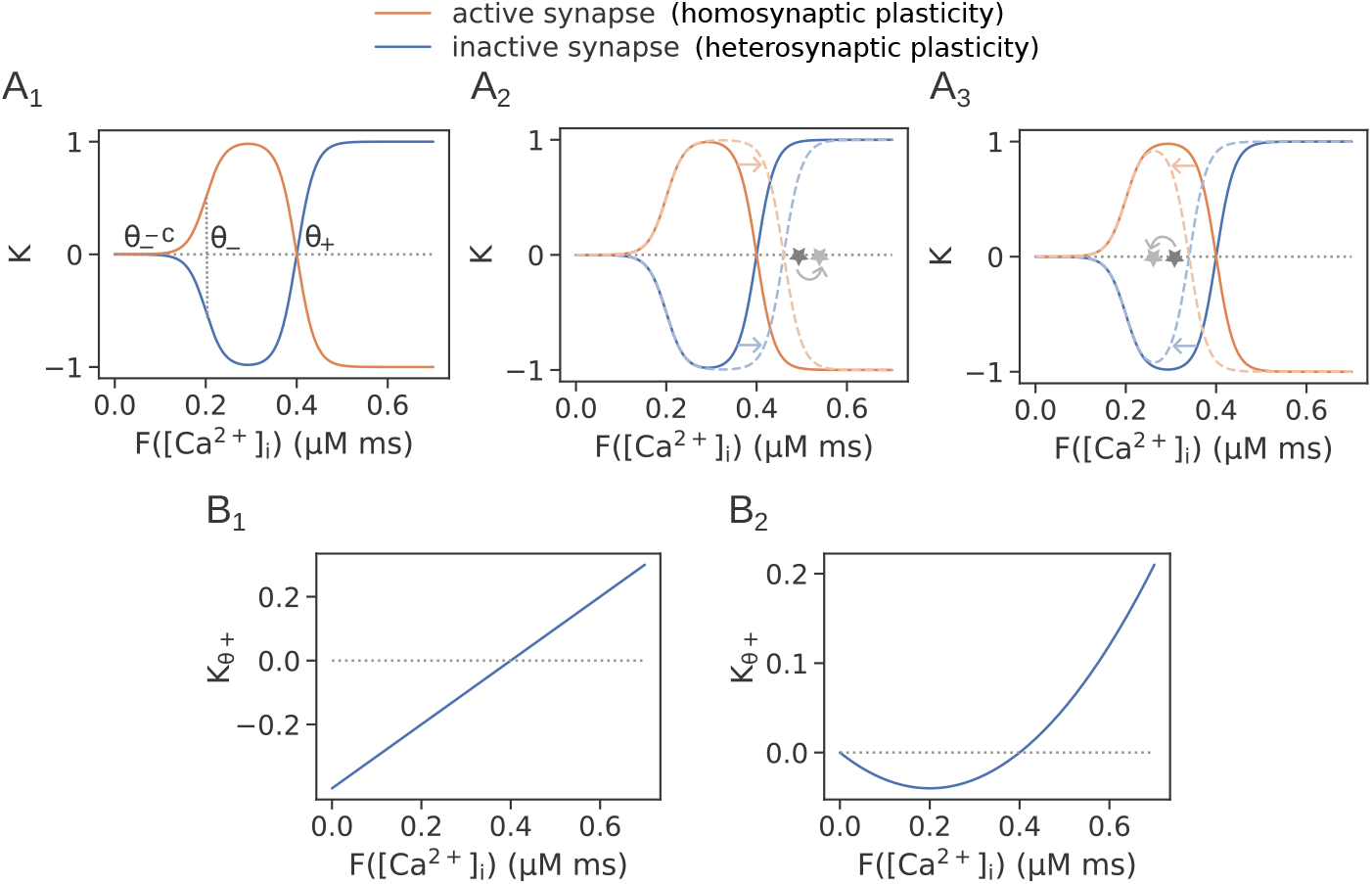
Kernels for homo- and heterosynaptic plasticity used in tandem in the second type of simulations. (A_1_) The plasticity kernels for active and inactive inhibitory synapses (homo- and heterosynaptic plasticity, respectively). The thresholds *θ*_+_ and *θ*_−_ −*c* are the same for both kernels. (A_2_, A_3_) Illustrations of how the plasticity kernels in (A_1_) change when *θ*_+_ is updated according to one of the metaplasticity kernels in (B_1_, B_2_). (A_2_) An update step when max (*F*) is greater than *θ*_+_. (A_3_) An update step when max (*F*) is smaller than *θ*_+_. Arrows indicate direction and amount of kernel movement during update. Solid lines depict kernels before the update, and dashed lines indicate kernels after the update. Dark grey star indicates max (*F*) before the update, and light grey star indicates max (*F*) after the update. (B_1_) The first variant of the metaplasticity kernel resulting in unstable weights with heterosynaptic plasticity (Eqs. 8 and 9). *θ*_+_ moves equally in each direction with this kernel. (B_2_) A second variant of the metaplasticity kernel used to ensure the stability of the inhibitory weights with heterosynaptic plasticity (Eq. 10). *θ*_+_ moves more easily towards higher values of *F* with this kernel. A nonlinear kernel such as this one is necessary for stability in the original BCM rule [11].

**Fig 10.**
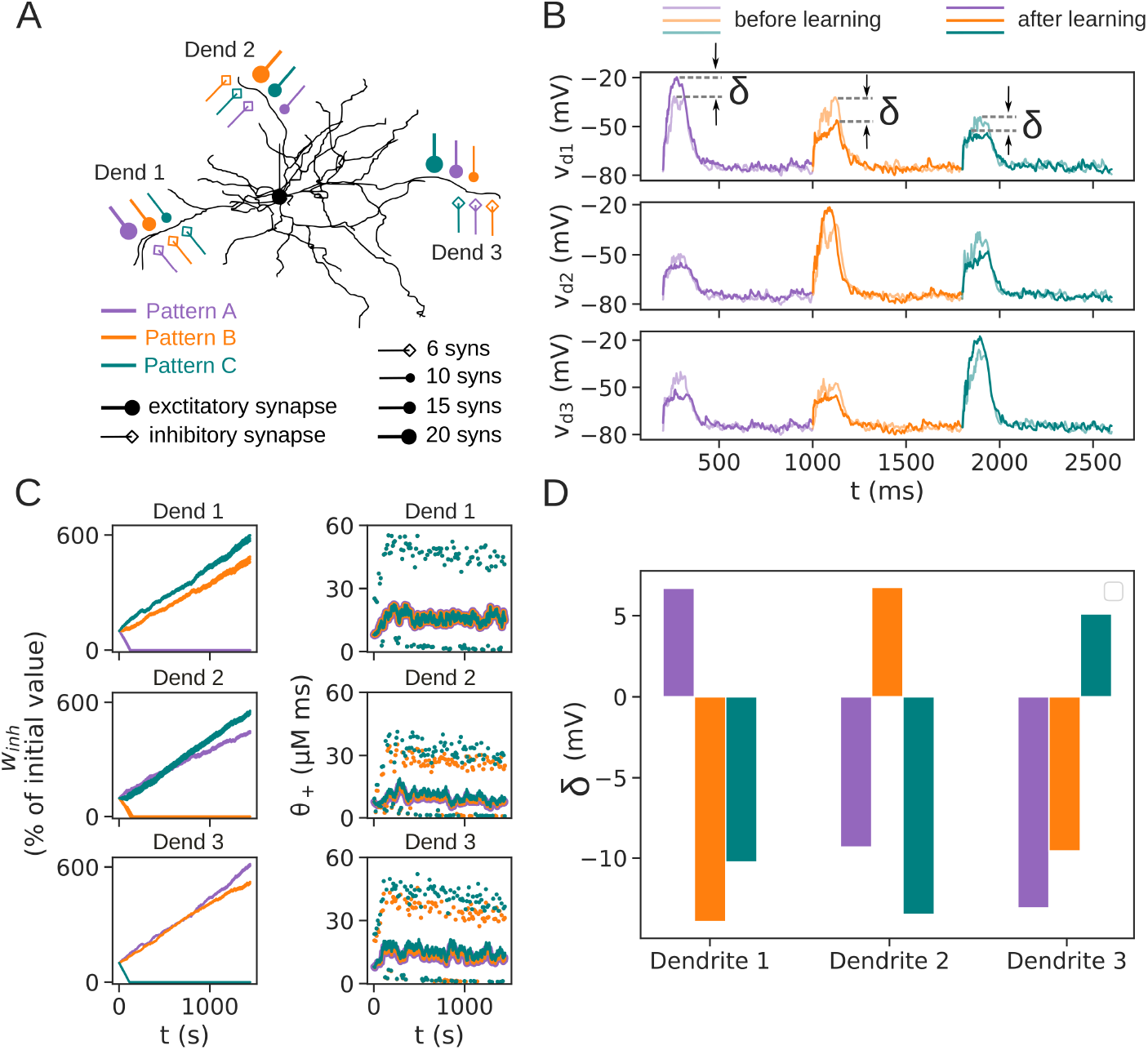
Control of supralinear dendritic integration with heterosynaptic inhibitory plasticity. Metplasticity occurs according to the same rule as in homosynaptic plasticity shown in Fig. 9B_1_ and given with Eq. 9 (where *θ*_+_ follows the maximal value of *F*). (A) The input configuration is the same as in Fig. 6. (B) Dendritic voltage before and after inhibitory synaptic plasticity. In each dendrite, the pattern with the largest excitiatory cluster evokes a stronger response after learning, while the two other patterns are inhibited. (C) (Left column) The evolution of inhibitory synaptic weights during learning. Inhibitory synapses for the strongest pattern in each dendrite are weakened, allowing it to evoke even stronger dendritic responses. Inhibitory synapses for the other two patterns are strengthened, inhibiting their dendritic responses. (Right column) The evolution of the threshold *θ*_+_ during learning (different line thickness is used to highlight overlapping plots). The maximal value of *F* during a pattern is shown as dots for the first inhibitory synapse in each cluster (overlapping dots are not highlighted). With this metaplasticity rule, the threhsold *θ*_+_ does not stabilize, and the weights grow without bounds. (D) The average change in dendritic depolarization after learning. In each dendrite, only the strongest pattern is allowed, while the other two patterns are inhibited. The same number of trials and averaging as in Fig. 8D is used.

**Fig 11.**
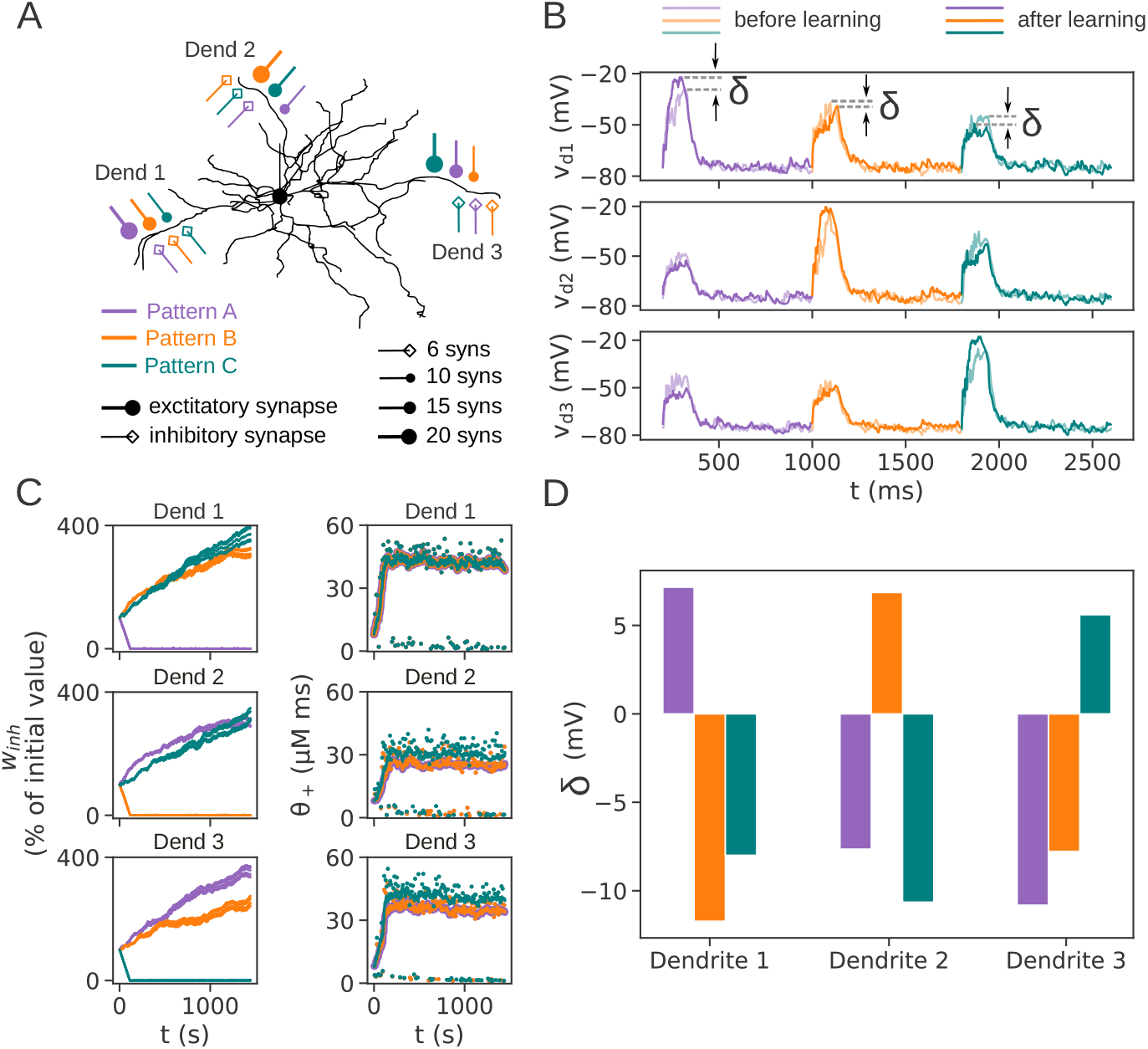
Control of supralinear dendritic integration with heterosynaptic inhibitory plasticity. Metplasticity is according to the second variant, shown in Fig. 9B_2_ and given with Eq. 10. (A) The input configuration is the same as in Fig. 8. (B) Dendritic voltage before and after inhibitory synaptic plasticity shows qualitatively the same result as in Fig. 10B: only the strongest pattern in each dendrite is allowed, and the other two patterns are inhibited. (C) (Left column) The inhibitory synaptic weights stabilize during learning compared to Fig. 10C. Inhibitory synapses for the strongest pattern in each dendrite are weakened, while those for the other two patterns are strengthened, as in Fig. 10C. (Right column) The evolution of the threshold *θ*_+_ during learning (different line thickness is used to highlight overlapping plots). The maximal value of *F* during a pattern is shown as dots for the first inhibitory synapse in each cluster (overlapping dots are not highlighted). (D) The average change in dendritic depolarization after learning. In each dendrite, only the strongest pattern is allowed, while the other two patterns are inhibited. The same number of trials and averaging as in Fig. 6D is used.

#### Heterosynaptic inhibitory plasticity speeds up learning to control supralinear dendritic integration and ensures stability of synaptic weights

We next examine what happens when heterosynaptic plasticity is added. We considered the following findings when constructing the rule for heterosynaptic plasticity. First, heterosynaptic plasticity can occur with or without the activation of nearby exctitatory synapses (i. e. elevation of the local voltage is enough), each with an opposite plasticity outcome [19]. Next, in the case when excitatory synapses are activated, they themselves often show plasticity together with the heterosynaptic inhibitory plasticity, which can be either of the same type (eLTP/iLTP and eLTD/iLTD; the prefixes “e” and “i” refer to excitatory and inhibitory, respectively) or the opposite type (eLTP/iLTD and eLTD/iLTP) [13, 14]. Inspired by this, we have constructed the kernel for heterosynaptic plasticity to be opposite (inverted) of that for homosynaptic plasticity (Fig. 9A_1_), and we test only the scenario for gating the supralinear dendritic responses. Figures 9A_2_, A_3_ illustrate a single update of the threshold *θ*_+_ when *F* is above or below it. Figure 9B_1_ shows the metaplasticity rule used with heterosynaptic plasticity, and Fig. 9B_2_ shows another variant of metaplasticity that ensures stability of the modified inhibitory synaptic weights.

With heterosynaptic plasticity, besides the homosynaptically-modified active inhibitory synapses belonging to one pattern, the inactive inhibitory synapses belonging to the other two patterns are also modified. The active synapses are modified according to the kernel for homosynaptic plasticity (Fig. 9A_1_, orange line), while the inactive synapses are modified according to the inverted kernel for heterosynaptic plasticity (Fig. 9A_1_, blue line). Values of *F* below the threshold *θ*_+_ (but above *θ*_−_), which correspond to low [Ca^2+^]_i_, trigger homosynaptic LTP in the active inhibitory synapses, and heterosynaptic LTD the inactive ones. Conversely, values of *F* above the threshold *θ*_+_, corresponding to high [Ca^2+^]_i_, elicit homosynaptic LTD in the active inhibitory synapses and heterosynaptic LTP in the inactive ones.

Adding heterosynaptic plasticty speeds up learning compared to homosynaptic plasticity only. This occurs because during one update step, both the active and inactive synapses are modified, meaning that synpases representing all patterns are modified, even though only one of the patterns is activated. Because the inactive synapses do not need to “wait” to be activated in order to be modified, it takes less pattern repetitions to achieve the same result as homosynaptic plasticity (Fig. 10C, left, synapses are weakened faster, after around 200 pattern presetntations, compared to Fig. 9C where more than 500 pattern presentations are required). Heterosynaptic plasticity results in only the strongest pattern to be allowed in each dendrite, while the other two patterns to be suppressed (Fig. 10). This is most clearly seen in the dendritic voltage responses to each pattern, before and after inhibitory synaptic plasticity (Fig. 10B). For example, in dendrite 1, only the strongest pattern A is allowed, while patterns B and C are suppressed, and the analogous outcome occurs in dendrites 2 and 3 (Fig. 10B). The inhibitory synaptic weights in the first column of Fig. 10C and the average dendritic depolarization over 128 trials in Fig. 10D show the same result: inhibitory synapses for the strongest pattern in each dendrite are weakened, while those for the other two patterns are strengthened. Comparing with the results for homosynaptic plasticity in Fig 8D, we note that even though the initial values of the threshold *θ*_+_ are the same in both scenarios, only the strongest pattern is disinhibited with heterosynaptic plasticity. This is a result of the kernel being opposite to that of homosynaptic plasticity: when the strongest pattern causes homosynaptic weakening of its inhibitory synapses, inhibition to the other two patterns is heterosynaptically increased. This will lower future supralinear responses of the other patterns, situating their calcium activity below the threshold when their own homosynaptic plasticity is triggered, inhibiting them further (even if before learning they may have evoked calcium activity above the threshold).

However, because the threshold *θ*_+_ for the same synapse is increased when a synapse is active and decreased when it is inactive, its weight never stabilizes in value (second column in Fig. 10C), causing the inhibitory weights to grow without bounds (weights are increased 6-fold in Fig. 10C, first column). This instability is removed with the second variant of metaplasticity, shown in Fig. 9B_2_, where the dependence of *θ*_+_ on *F* is nonlinear (quadratic) (Fig. 11). Again, only the strongest pattern is allowed in each dendrite, and the other two are suppressed (Fig. 11B). However, the inhibitory synaptic weights are not increased as much, and their levels stabilize (only around 4-fold increase in Fig. 11C, first column, compared to Fig. 10C). Figure 11C also shows that the sliding threshold *θ*_+_stabilizes in value, resulting in stable synaptic weights. Repeating this experiment for 128 trials also shows the same average change in dendritic depolarization as in the first metaplasticity variant in heterosynaptic plasticity (Fig. 11D). Moreover, heterosynaptic plasticity also resolves the weight instability that appeared in homosynaptic plasticity for the strongest pattern, because it always decreases the threshold *θ*_+_ by a small amount when a synapse is inactive, countering the increase made when the synapse is active. This keeps the threshold within small region below *F*, preventing any variations in *F* to trigger the positive feedback loop initiated when only homosynaptic plasticity is used (Fig. 11C, right column).

Lastly, it did not make sense to examine the effect of heterosynaptic plasticity with an inverted kernel in the scenario for equalizing the dendritic response amplitude. This is because the inverted kernel would make heterosynaptic plasticity act as a positive feedback loop, pushing the evoked voltage away from the set point, and counteracting the effect of the negative feedback loop from homosynaptic plasticity. Another option in the scenario for equalizing the dendritic responses could have been that both homosynaptic and heterosynaptic plasticity use the same kernel in Fig. 3. This also does not make sense because, for example, when the largest cluster is activated. the evoked value of *F* will be above the threshold in all nearby synapses, including the inactive ones. This will trigger homosynaptic plasticity in the correct direction for the active synapses, but heterosynaptic plasticity for the inactive synapses will be in the opposite direction of that needed for equalizing the dendritic responses. For this reason, we did not explore heterosynaptic plasticity in this scenario.

#### Control of supralinear dendritic integration can support nonlinear feature binding

Monitoring changes in inhibitory synapses across the whole dendritic arbor of cortical pyramidal L2/3 neurons has shown that such changes cluster together with changes in dendritic spines, many of which possibly represent excitatory synapses [20]. This example, together with the many results reviewed in [14], suggests that inhibitory synaptic plasticity occurs in concert with excitatory synaptic plasticity to shape neuronal output. Therefore, in this section we demonstrate that the control and gating of supralinear dendritic excitations also works in a more dynamic scenario where the excitatory synaptic clusters are not fixed, but are also plastic. For this we use a simple excitatory plasticity rule for corticostriatal synapses onto dSPNs, explained below and in the Methods section [21]. In particular, we show that inhibitory synaptic plasticity can support solving the nonlinear feature binding problem (NFBP) by a single SPN. The NFBP is a computationally difficult task because it belongs to the class of linearly non-separable problems, whose solution requires networks of artificial computational units (a single unit cannot solve it). In its simplest form, the NFBP consists of discriminating between two types of patterns (or stimuli), each of which is a combination of two features (Fig. 11A). It is most commonly illustrated with examples from visual feature binding, although it is potentially relevant to all brain regions which perform integration of multimodal inputs, or inputs representing different features of the same modality [22–24]. To solve the NFBP, the neuron should respond with somatic spiking to only two patterns (here called relevant patterns) while remaining silent for the other two (irrelevant patterns). The problem is nonlinear since, on average, the neuron receives the same amount of excitation for all four patterns, but should nevertheless learn to respond to these patterns differently.

To illustrate how inhibition can solve the NFBP, we use the setup in Fig. 11B. This was also done in [17], so in this section and the next one we extend the investigation to all possible setups (input configurations) of four features on two dendrites. In the input configuration in Fig. 11B, three features are present with excitatory clusters of 10 synapses in two dendrites, such that both relevant patterns can be represented by the same neuron, each in a different dendrite. The features are shared between the relevant and irrelevant patterns (for example, the feature ‘banana’ is shared by both the ‘yellow banana’ and ‘red banana’ pattern on dendrite 1). Because the excitatory clusters are equal in size, each pattern activates 20 excitatory synapses. This means that, initially, no pattern is stronger than the other, and inhibitory plasticity has no way of choosing which pattern to suppress in the dendrite (as it could in the simulations in the previous two sections, due to the different strength of the patterns). The stimulation protocol consists of presenting patterns sequentially, after which dopaminergic feedback, which guides excitatory synaptic plasticity, is delivered (Fig. 11C). A dopamine peak, which gates LTP in the excitatory synapses, is delivered for the relevant patterns, and a dopamine pause, which gates LTD, is delivered for the irrelevant patterns. The excitatory plasticity rule is very simple, increasing or decreasing weights towards pre-specified maximal and minimal values. Nevertheless, it is biologically-grounded in the sense that plasticity is triggered by signals relevant for the corticostriatal synapse onto dSPNs (see Figure 4 - figure supplement 2 in [21] and the Methods section for details). The activity of inhibitory synaptic clusters is again longer than that of the excititatory pattern, as it needs to suppress long-lasting supralinear dendritic responses. In addition, theoretical considerations have suggested that the supralinear dendritic voltage elevation should be all-or-none in amplitude to solve the NFBP. Because of this, we have used plateau potentials in this scenario, triggered when 20 excitatory synapses become sufficiently strong (using glutamate spillover; see Methods for details and [25]).

Excitatory synaptic plasticity causes strengthening of the synaptic clusters which represent relevant patterns and weakening of the remaining cluster taking part in the irrelevant pattern (Fig. 11F, upper panels). This causes the emergence of a ‘stronger’ pattern in each dendrite, which is then “used” by the inhibitory plasticity rule. As in the previous example with fixed synaptic weights, inhibitory synapses corresponding to the stronger excitatory stimulus weaken, while those for the other stimulus strengthen, eventually learning to gate only one relevant pattern in each dendrite (Fig. 11F, lower panels). The excitatory plasticity alone cannot solve the NFBP. This is because the simple excitatory plasticity rule always strengthens weights to their maximal value (Fig. 11F, upper panels), i.e. it does not know how to stop the weights at a value where only the relevant patterns will trigger supralinear dendritic responses. This is exemplified with the dendritic and somatic voltage traces in Fig. 11E, which show that after learning, both relevant and irrelevant patterns elicit somatic spiking. The reason why the irrelevant patterns also trigger supralinear dendritic responses is that half of their excitatory synapses (those shared with a relevant pattern) have been strengthened to their maximal values. With the inhibitory plasticity rule, inhibitory synapses learn to suppress the supralinear dendritic responses for the irrelevant patterns, gating through only the supralinear responses for the relevant patterns. In this way, somatic spiking is elicited only for the relevant patterns, as can be seen from the voltage traces in Fig. 11D. In addition to the input configuration in Fig. 11B, we also ran the learning simulation on all possible 18 input configurations that can solve the NFBP (shown in S3 Fig), verifying that the NFBP could not be solved with excitatory plasticity alone - inhibitory plasticity was also needed (Fig. 11G, see Methods for a description of the performance measure). In hippocampal CA1 pyramidal neurons, inhibitory synapses from interneurons have been found to prevent supralinear dendritic integration which drives burst spiking in the soma [9]. This is also what the learning rule for the inhibitory synapses achieves in the NFBP when inhibition prevents the supralinear dendritic responses for the irrelevant patterns, leaving burst spiking evoked only by the relevant patterns.

Lastly, we note that if the supralinear dendritic excitations evoked by the relevant patterns are large and all-or-none in their voltage amplitude (like the plateau potentials generated with glutamate spillover in this study), there are strong indications that a different excitatory learning rule alone (which “knows” when to stop strengthening the synapses) is enough to solve the NFBP [21]. Without such an excitatory plasticity rule, as is the case in this study, inhibition is necessary. Also, if the supralinear dendritic responses are not all-or-none, i.e. are graded in amplitude, then again a combination of excitatory and inhibitory plasticity is necessary to solve the NFBP [17].

#### Targeted inhibition can solve the NFBP in difficult cases

The NFBP is difficult to solve when all four features innervate one dendrite. Figure 11A shows such a “symmetric” case with two layers of symmetry. The first layer consists in the fact that before learning, a dendrite’s response is undifferentiated with respect to all four patterns, i.e. it responds equally to all patterns. During learning, it needs to pick one of the relevant patterns to strengthen. The second layer consists in the fact that each dendrite needs to store a different pattern, despite both dendrites being initially undifferentiated with respect to all patterns. In situations such as this one, excitatory plasticity alone is on average not enough to make the differentiation and solve the NFBP - usually the same pattern is stored in both dendrites. This is shown in Fig. 11D, where two plateau potentials are evoked for the yellow banana, one in each dendrite, which are a result of the same synapses being strengthened in both dendrites (Fig. 11F). Only the first layer of symmetry is broken, and which pattern is stored is determined by the random sequence of arriving patterns. A different mechanism, such as e.g. branch plasticity, which modifies the strength of signal propagation through the dendrite, has been proposed for breaking the second layer of symmetry in these cases [26].

Here we show that targeted inhibition on the dendrites can solve the NFBP in cases requiring symmetry breaking. For example, in the case in Fig. 11A, each dendrite receives inhibitory synapses meant to inhibit specific features. These synapses break the first layer of symmetry in each dendrite - they determine which pattern will be stored by inhibiting the features belonging to the other pattern. Through inhibition, a “stronger” pattern is created in each dendrite, which excitatory plasticity strengthens further and stores in the dendrite (Fig. 11E). If inhibition targets different features in the two dendrites dendrites, as in Fig. 11A, it can also break the second layer of symmetry. This is seen in Fig. 11E, where dendrite 1 stores the red strawberry, and dendrite 2 stores the yellow banana. Each pattern evokes a plateau potential in its dendrite after learning, causing somatic spiking, while no plateaus nor somatic spikes are evoked for the irrelevant patterns (Fig. 11C).

We have tested all possible symmetric arrangements of four features on two dendrites, shown in S4 Fig. In each of these configurations, all foru features innervate at least one dendrite. Also, the dendrites receive targeted inhibitory synapses which suppress dendritic activity from the features that need to be weakened. When inhibition is pre-arranged in this way, inhibitory plasticity enables the dendrites to become responsive to only one pattern - the one which is not inhibited. Thus, inhibition is the differentiating (symmetry-breaking) factor that allows the NFBP to be solved in the symmetric cases (Fig. 11B, red trace). Without it, the same pattern is stored in both dendrites on average, and the NFBP is not solved (Fig. 11B, yellow trace).

**Fig 11.**
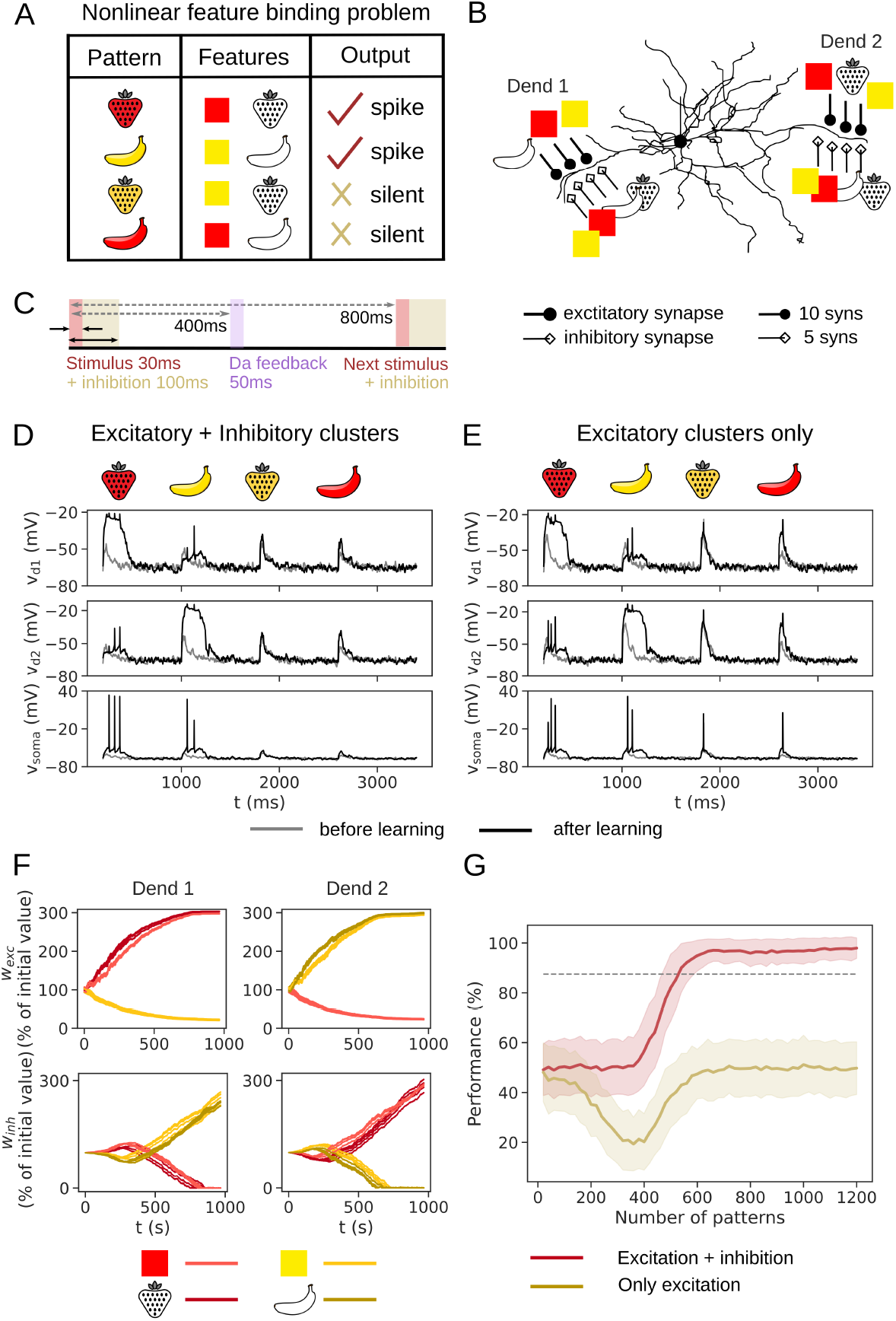
Control of supralinear dendritic responses used to solve the NFBP (nonlinear feature binding problem). (A) An example of the NFBP from the visual system. A pattern constists of two features (color and shape), each of which can have two values (‘red’ and ‘yellow’, and ‘strawberry’ and ‘banana’), giving a total of four possible patterns. In order to solve the NFBP, a neuron should spike to only two patterns (feature combinations) and remain silent for the other two. (B) The input configuration used in these simulations. The features are represented with both excitatory and inhibitory synaptic clusters of 10 and 5 synapses, respectively. Excitatory synaptic clusters for only three features arive on one dendrite, together representing one relevant and one irrelevant pattern. The two relevant patterns are represented in different dendrites. Conversely, inhibitory synaptic clusters for all four features innervate both dendrites. (C) The stimulation protocol in used in the learning trials. Excitatory synaptic clusters are activated in a window of 30 ms with 3 spikes per synapse, while inhibitory synaptic spikes are activated in a window of 100 ms with 10 spikes per synapse, both windows having the same starting time (the inhibition window is longer in order for effective suppression of supralinear dendritic responses). After 400 ms the dopamine feedback is delivered, whereas the next stimulus arrives after 800 ms. (D) Voltage traces from the dendrites and soma for each pattern before and after learning, for the input configuration in (C) containing both excitatory and inhibitory synapses. (E) The same plot as in (D) when only excitatory synapses are present. (F) Evolution of excitatory and inhibitory synaptic weights in the two dendrites for the input configuration in (C). Performance on the NFBP for the scenario with and without inhibitory synaptic clusters. Results are averages over 18 possible input configurations that can solve the NFBP where up to three features innervate one dendrite (S3 Fig), with 13 trials per one input configuration (for a total of 234 trials). In each trial, a pair of dendrites is randomly chosen from a set of 9 dendrites, and 1200 pattern presentations are used.

**Fig 11.**
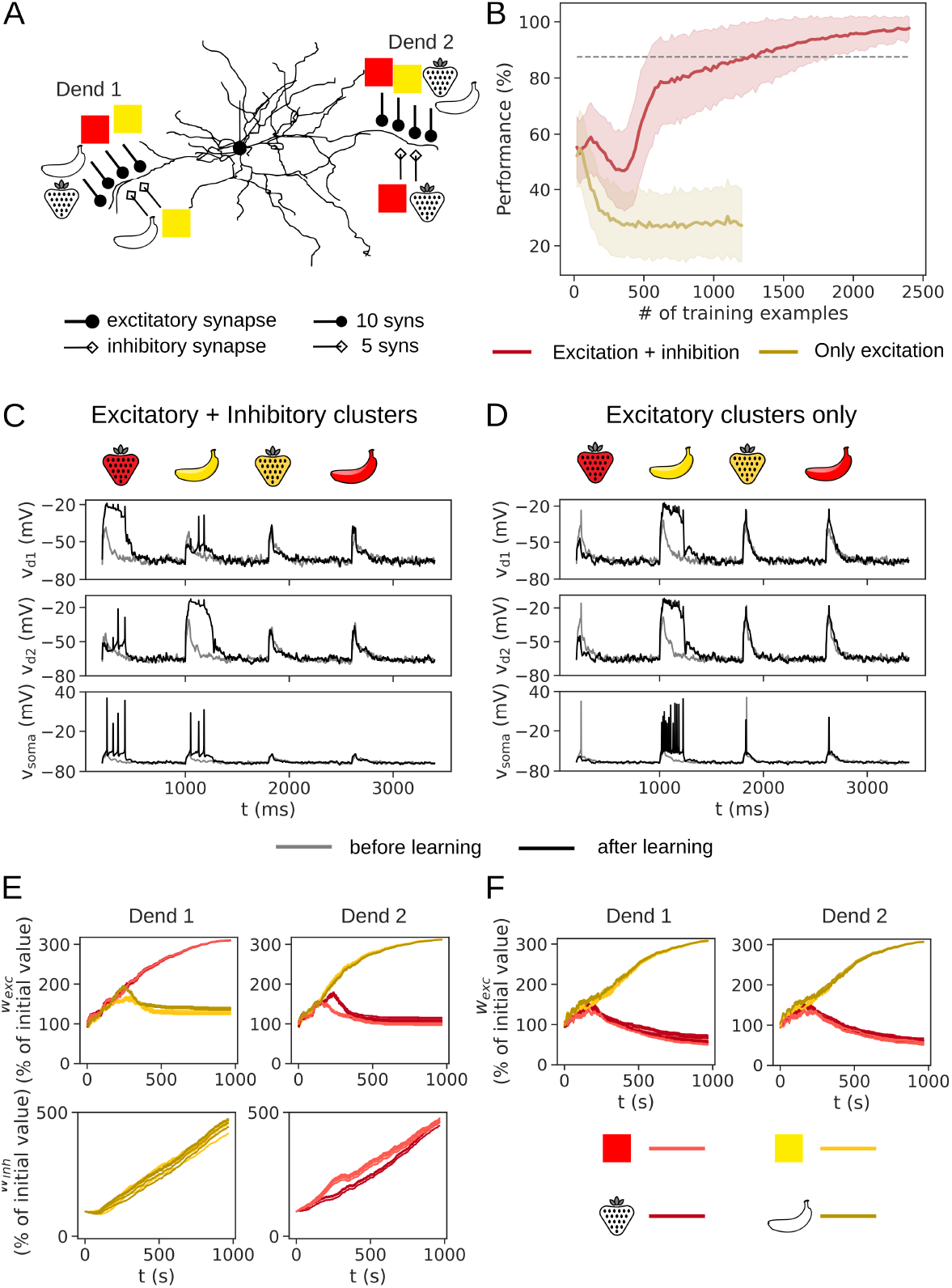
Targeted inhibition can solve the NFBP in difficult (symmetric) cases where all four features innervate at least one dendrite. (A) An example of such a difficult input configuration with all four features present on two dendrites. Inhibitory synapses only for certain features innervate the dendrites. (B) Performance on the NFBP averaged across all input configurations with four features on at least one dendrite, shown in S4 Fig. 19 trials are performed for each input configuration, for a total of 247 trials. In each trial, synapses are placed on two randomly chosen dendrites from a set of 9 dendrites. When inhibition targets only certain features, the NFBP can be solved. Otherwise, on average the same pattern is stored in both dendrites without inhibition, and the NFBP is not solved. In the latter case the learning was run for a twice shorter amount of time in order to save computational resources. (C) Voltage traces from the dendrites and soma for each pattern before and after learning, for the input configuration in (A) containing both excitatory and inhibitory synapses. (D) The same plot as in (C) when only excitatory synapses are present. (E) Evolution of excitatory and inhibitory synaptic weights in the two dendrites for the input configuration in (A). (F) Evolution of excitatory synaptic weights when no inhibitory synapses are present in the input configuration in (A).

## Discussion

In this study we demonstrate a local inhibitory plasticity rule (or a set of related rules) that can control the neuronal firing rate during continuous, distributed, rate-coded input signals, and can control supralinear integration in dendrites during sparsely-coded, clustered inputs. The rule is based on the BCM formalism, and updates weights based on a filtered variable of local dendritic L-type [Ca^2+^]_i_. For rate-coded inputs, the results show that altered neuronal firing rate caused by changes in excitatory input rate or excitatory synaptic strength can be completely compensated by inhibitory synaptic plasticity, restoring the original neuronal firing rate before the change (as long as there is enough excitation for the neuron to spike). This balance of the total neuronal excitatory drive with a matching inhibitory drive is a well-established function of inhibitory synapses, and one of its roles is to make neurons responsive to a wide range of excitatory input frequencies [2–5, 27, 28].

For sparsely-coded inputs, the rule enables control of supralinear dendritic integration in two different modes when several patterns are represented in a single dendrite, depending on the function (kernel) that translates calcium activity to changes in weight. In one mode, the rule can equalize dendritic responses arising from excitatory clusters of different sizes. This makes a dendrite (and the neuron) responsive to which patterns are active, not how strong they are. This normalization of pattern (or feature) strength on the level of dendrites is similar to the normalization of individual synapses so that they contribute equally to somatic excitation regardless of their distance, a term called synaptic democracy [29]. Thus, in this mode, the inhibitory plasticity rule enables a similar “cluster democracy” at the level of a dendrite (or at least at the level of a dendritic region containing nearby clusters). In the other mode, the rule can learn to permit or inhibit supralinear dendritic responses in a dendrite. Here, homosynaptic plasticity alone results in unstable synaptic weights, and heterosynaptic plasticity provides both weight stability and faster learning (requires less pattern presentations). Gating supralinear dendritic integration is another well-established role of inhibitory synapses [6–9]. Moreover, changes in inhibitory synapses have been found to cluster with changes in dendritic spines in L2/3 pyramidal cortical neurons, suggesting that modifications in their synaptic weights can at least regulate local dendritic activity [20].

These results demonstrate that if inhibitory synapses whose plasticity is governed by the BCM rule exist, they would be versatile enough to enable learning how to execute two important functions of inhibition: i) control of the neuronal firing rate, and with it, extending a neuron’s dynamical range, and ii) gating or inhibiting supralinear dendritic responses, just by using signals available locally at the synaptic site. Furthermore, the rule we have demonstrated is flexible in the sense that other factors, such as e.g. neuromodulation, could decide whether homo- or heterosynaptic inhibitory plasticity is engaged, as well as which of the two opposite kernels for plasticity is used.

A significant number of inhibitory synapses has been found to be on spines that receive excitatory inputs [20]. We have not explored this scenario in the present study, but inhibitory synapses targeting single spines have been shown to control [Ca^2+^]_i_ levels with single spine resolution [30]. As such, inhibitory plasticity could regulate excitatory synaptic plasticity in a single spine, affecting that spine’s future participation in synaptic clusters.

### Plausibility of the inhibitory plasticity rule

The molecular mechanisms underlying inhibitory synaptic plasticity are known in greatest detail for the inhibitory synapses ariving on Purkinje neurons; specifically, they detail the postsynaptic biochemical machinery responsible for LTP [16, 31–34]. In these synapses, strong depolarization alone without any presynaptic activation (i.e. when synapses are inactive) causes LTP, which is evoked by high calcium influx. In the postsynaptic molecular circuit, both simulations and experiments have suggested that the amount of phosphodiesterase 1 (PDE1) determines the calcium threshold for eliciting LTP: more PDE1 causes a higher threshold, and less PDE1 causes a lower one [16]. The existence of a threshold and the occurence of LTP for high calcium influx fits the kernel for heterosynaptic inhibitory plasticity in Fig. 9A_1_, and if a mechanism for upregulating the amount of PDE1 were discovered after high activity, this would fit the sliding threshold in the kernel. However, despite the known fast regulation of PDEs by calmodulin, kinases, and anchoring proteins, mechanisms for longer-term regulation of PDE amounts, which in turn regulate the calcium threshold for plasticity, seem to have not been identified yet, at least in neurons [35, 36]. Nevertheless, anchoring proteins, which organize postsynaptic signalling and which also bind PDEs, are subject to activity-dependent regulation [37]. Because of this, it is possible they are involved in a mechanism that implements a sliding calcium threshold through their interactions with PDEs. In passing, we note that the molecular circuitry in the inhibitory synapses to Purkinje neurons studied in [16] is also present in the excitatory synapses in the striatum, where instead of PDE1, PDE10A is predominantly expressed, and anchored to AKAP150 [38, 39]. This raises a possibility that PDE10A controls the calcium threshold in excitatory synaptic plastiticy.

On the other hand, if the inhibitory synapses on Purkinje neurons are activated together with the applied strong postsynaptic activity (thus also also lowering calcium influx), then no LTP occurs [16, 34]. However, an analogous, postsynaptic, LTD-triggering molecular circuit has not been estabilshed in these neurons, so it is not possible to discuss further whether lower calcium influx causes LTD and can be related to the BCM-type kernel in this study.

Many pieces of experimental evidence exist that show that excitatory and inhibitory plasticity interact, and that plasticity in one synaptic type, e.g. the excitatory synapses, triggers heterosynaptic changes in the other synaptic type, e.g. the inhibitory synapses [14]. Very often, these changes are in the opposite direction, i.e. eLTP occurs together with iLTD, and eLTD with iLTP [14]. With the inhibitory plasticity rule in this study, this outcome occurs in the simulations for the NFBP, where features whose excitatory synapses are strengthened have their corresponding inhibitory synapses weakened, and vice versa. Sometimes, the plasticity outcomes in excitatory and inhibitory synapses are in the same direction (eLTP/iLTP and eLTD/iLTD) [14]. This occurs in the simulations for controlling the neuronal firing rate. However, in both of these simulations, the inhibitory plasticity rule is homosynaptically induced, i.e. it requires that inhibitory synapses are activated, while in the experiments it is often heterosynaptically induced, through molecular crosstalk between excitatory and inhibitory plasticity. We have not implemented such crosstalk explicitly (apart from using intracellular calcium concentration evoked by the local voltage). Instead, it is the shape of the inhibitory plasticity kernel that determines whether inhibitory plasticity will occur in the same or the opposite direction of excitatory plasticity.

### Collateral SPN-to-SPN synapses could contribute to solving nonlinear feature binding

We have demonstrated the use of controling supralinear integration in solving a linearly non-separable task, the NFBP, where inhibitory plasticity works together with a simple excitatory plasticity rule that describes plasticity in corticostriatal synapses onto dSPNs. In the following we describe why inhibition could be particularly relevant for solving this task in the striatum. The striatum is the input nucleus of the basal ganglia and is thought to be involved in selecting actions (movements), or initiating actions already selected by the cortex [40, 41]. The two different types of SPNs are part of two basal ganglia pathways - the direct pathway, which initiates movements, and the indirect pathway, which suppresses movements, and the balance of activity between these two pathways is important for proper movement initiation (Fig. 12) [42, 43]. The SPNs also send collateral inhibitory projections to other SPNs, which could play a role in achieving this balance [44]. In the NFBP, the relevant patterns, such as the red strawberry, need to increase the activity of the direct pathway and reduce that of the indirect pathway. Conversely, the irrelevant patterns, such as the red banana, need to inactivate the direct pathway and activate the indirect pathway. In this study we only exemplified how direct-pathway SPNs (dSPNs) can learn to respond to relevant patterns with a combination of excitatiory and inhibitory synaptic plasticity. However, the indirect-pathway SPNs (iSPNs) have an analogous excitatory synaptic plasticity mechanism with which they can learn to respond to the irrelevant patterns (in order to suppress actions for these patterns) [45–47]. Because the patterns in the NFBP share features (such as ‘red’ in the red strawberry and red banana), collateral inhibitory synapses between dSPNs and iSPNs could have a role in inhibiting the other pathway’s response to the shared feature (Figs. 12B and 12C). For example, imagine that the red strawberry is stored in synaptic clusters to the dSPNs, and that the red banana is stored in synaptic clusters to the iSPNs. When each of them arrives, it would increase the activity of the respective pathway (green highlight in Fig. 12B and purple highlight in Fig. 12C). At the same time, through the collateral SPN projections, each of them can simultaneously inhibit the activity of the opposite pathway, which is evoked by the shared ‘red’ feature (highlighted collateral inhibitory projection decreases activity in the opposite pathway in Figs. 12B and 12C). Indeed, SPN-to-SPN projections form inhibitory inhibitory synapses that are located on the dendrites, and are thus suitable for affecting dendritic integration [48]. The inhibitory plasticity rule that we presented is activated by local calcium levels evoked by excitatory synapses. Because of this, as excitatory plasticity develops, inhibitory plasticity will follow and match the strength of collateral inhibitory projections, which is suitable for developing the connectivity to solve the NFBP.

**Fig 12.**
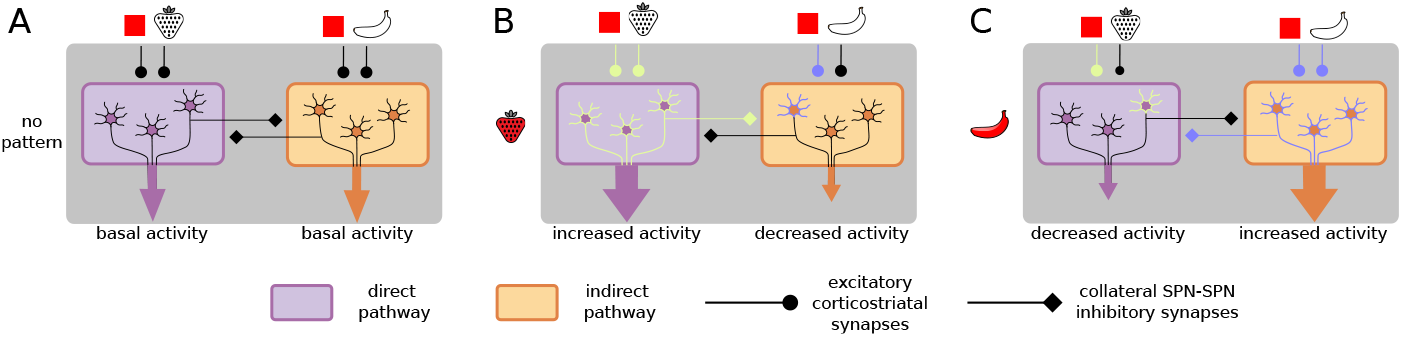
The possible role of collateral inhibition between SPNs of the direct and indirect pathway in solving the NFBP. (A) No pattern is presented, and the activity of the two pathways is balanced at a basal level. (B) A relevant pattern arrives (here the red strawberry) activating the clusters for the features ‘red’ and ‘strawberry’ on the dSPNs which have learned to respond to this pattern. It also activates the clusters for ‘red’ in the iSPNs which have learned to respond to the red banana. The increased activity of the direct patwhay can inhibit the iSPNs activated by the feature ‘red’, which could otherwise suppress movement. (C) The analogous situation for the irrelevant pattern red banana. It increases the activity of the indirect pathway by activating the iSPNs which have learned to respond to this pattern. Activation of the indirect pathway could inhibit the direct pathway through collateral inhibition.

### Relation to other studies

The first function, control of the neuronal firing rate, has been achieved with a non-local inhibitory plasticity rule which uses the relative timing of incoming presynaptic spikes and emitted somatic spikes [49]. The timing of somatic spiking, which is a global neuronal signal, is not always available to all synapses, such as the distal synapses where backpropagating spikes may not reach. Therefore it remains a question how suitable and applicable such non-local learning rules are for distal synapses. In comparison, the local learning rule studied here, which enables learning by using only local calcium activity, is a step toward greater biological detail.

The spike-timing-dependent inhibitory rule in [49] is formulated so that the neuronal activity is compared to a predetermined neuronal firing rate (set point), and weights are updated based on that difference. A similar formulation is used in [50] weight changes are driven by the difference in local voltage and a set point (target) voltage. In our case, the kernel used for controlling the neuronal firing rate also implements a similar negative feedback loop based on the difference of the calcium activity from the kernel’s threshold (set point). Homeostatic changes occur after prolonged changes in input or neuronal activity, and are in general thought to occur through molecular mechanisms different from those of synaptic plasticity (even though the pathways may partly overlap and converge on the same effectors) [51]. Compared to homeostatic plasticity which may take days to take effect, inhibitory synaptic plasticity operates on a shorter timescale (minutes to hours). Nevertheless, homeostatic roles of inhibitory plasticity have also been reported [52].

Inhibitory plasticity acts in concert with excitatory plasticity to shape and fine-tune the responses of entire neural circuits as cortices develop and mature, giving rise to a so-called balance of excitation and inhibition (E/I balance), which has been studied extensively [53, 54]. How local learning rules create the so-called detailed and tight E/I balance both in recurrent neural networks and in individual neurons is not known [53, 54]. The local inhibitory rule studied here provide some answers to this question on the single neuron level, and could provide some on the network level if used in biophysically detailed neural circuit models [55, 56]. Also, it has been shown both with simulations and experiments that the learning rates of excitation and inhibition determine the final level of E/I balance in single neurons [57]. This is an additional way of regulating the outcome of inhibitory plasticity that we did not study and that can be addressed in the future.

## Methods

### Neuron model

We use a multicompartment model of a striatal projection neuron (SPN), taken from the collection of models published in [58]. Briefly, the neuron models in this collection are fitted to match several relevant electrophysiological properties by optimizing the conductances of ion channels expressed by SPNs, and are validated against experimental data for dendritic calcium responses evoked by backpropagating action potentials and data for somatic current-frequency responses. All details are described in [58].

Clustered excitatory synapses were placed on spines (as in [25]). Each spine was modeled as two additional compartments consisting of a neck and a head region, and contains voltage-gated calcium channels of types L (Ca_v_1.2 and Ca_v_1.3), R (Ca_v_2.3), and T (Ca_v_3.2 and Ca_v_3.3). The voltage-gated calcium channel conductances in the spines were also manually tuned to match the relative calcium proportions in [59] and [60], as well as the calcium amplitudes due to stimulation with backpropagating action potentials in [61]. The addition of explicit spines did not change the basic behavior of the model, such as the response to current injections, etc. Inhibitory synapses were not placed on spines.

### Synaptic inputs

#### Dual exponetial synaptic model (difference of two exponentials)

In the first type of simulations, excitatory and inhibitory synapses are distributed across the dendrites according to the synapse densities reported in [62], and they are not located on spines. The conductances of both excitatory and inhibitory synapses are modeled with a difference of two exponential functions, also called the dual exponential synaptic model. For a spike arriving at time 0 the synaptic conductance in this model is given by:

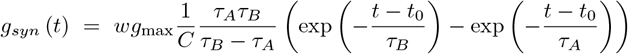

where *g*_max_ is the maximal synaptic conductance, and *τ*_*A*_ and *τ*_*B*_ are the rise and decay time constants. *C* is a normalization factor given with:

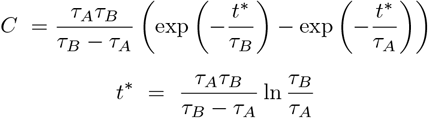

where *t*^*\**^ is the position of the maximum of the dual exponential function. This normalization ensures that the time-varying part of the dual exponential reaches a peak value of 1, which is then scaled by the maximal conductance *g*_max_ which has units of nanosiemens (nS). The synaptic weight, *w*, is a non-negative, dimensionless parameter which scales *g*_max_. It is modified by synaptic plasticity, thus changing the synaptic conductance. Here we also use the name synaptic strength for the product of the weight and the maximal conductance *wg*_max_, which has units of nS, to differentiate it from the dimensionless synaptic weight *w*.

The synaptic current arising from the conductance *g*_*syn*_ is:

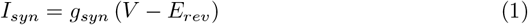

(In NEURON the model is implemented by a scheme integrating two first order differential equations which allows for convenient integration of spikes arriving at any time.)

Excitatory synapses have both AMPA and NMDA components, and NMDA synapses have a Mg^2+^ block described by a sigmoidal function:

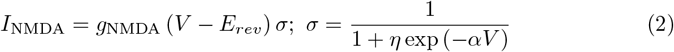

with parameters *η* = 0.38 and *α* = 0.062. The parameters of the dual exponential synaptic models for the AMPA, NMDA and inhibitory synapses are given in Table 1. Because our multicompartment neuron model contains less electrical compartments than the number of synapses reported in [62], this implies that several synapses will arrive in a single compartment. Since their synaptic inputs will be integrated in the same voltage variable, we have represented such multiple synapses with a single synapse instead, whose input frequency is scaled by the number of synapses it represents. This is why we have used a non-saturating synapse model for these synapses.

**Table 1.** The synapse parameters in the dual exponential synaptic model, used in the simulations for controlling the neuronal firing rate.

|  | AMPA | NMDA | inhibitory |
| --- | --- | --- | --- |
| $\tau_A$ (ms) | 1.9 | 2.76 | 0.1 |
| $\tau_B$ (ms) | 4.8 | 115.5 | 10 |
| $E_{\text{rev}}$ (mV) | 0 | 0 | -60 |
| $g_{\text{max}}$ (nS) | 0.1 | 0.1 | 1.5 |

#### Saturating synapse models

In the second type of simulations, excitatory and inhibitory synapses comprising the background synaptic noise are modeled with the dual exponential model described above, with the parameters given in Table 1. In contrast, the clustered excitatory synapses, located only in particular dendritic branches, are represented with saturating synapse models taken from [63], which are a variation of the saturating synapse models in [64]. We chose saturating synapse models in this case since the synaptic clusters receive temporally coincident inputs comprised of three spikes each, and we wanted to avoid single excitatory synapses reaching unrealistically high conductance values.

The saturating synapse models are kinetic models which operate according to two different kinetic schemes depending on the presence of neurotransmitters. When neurotransmitters arrive at the postsynaptic site, receptor dynamics are described by a kinetic scheme that models switching between closed (*C*) and open (*O*) states with rates of *α* and *β*:

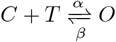

where *T* is the neurotransmitter. After the neurotransmitter has been cleared from the synaptic cleft, receptor dynamics evolves according to a kinetic scheme that describes just the closing of the receptor with a rate *β*:

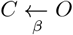

When a presynaptic spike arrives, neurotransmitter levels are assumed to always reach a fixed saturating concentration, *T*_max_, in the synaptic cleft, i.e., are represented by a pulse with amplitude *T*_max_ and duration of *T*_dur_. This “short pulse” representation of glutamate concentration in the synaptic cleft follows from findings in cultured hippocampal synapses showing that glutamate has a very fast time course in the cleft, rapidly reaching concentrations of 1 mM, with a decay constant of 1.2 ms [65]. If another presynaptic spike arrives while the neurotransmitter pulse is still “on”, the pulse duration is lengthened by *T*_dur_. In this model, the synaptic conductance is represented by the fraction of glutamate receptors in the open state at a given time:

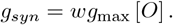

(The synaptic conductance is used to calculate the synaptic current according to Eq. 1.) This model uses the same rate of receptor closing irrespectively of whether transmitter is bound to the receptor or not, which is a simplification of biological reality, but still captures the waveform of the synaptic potential.

### Inhibitory plasticity rule

#### Indicator of synaptic activity

As an indicator of excitatory synaptic activity we use the level of intracellular calcium, [Ca^2+^]_i_, in the dendritic shaft where an inhibitory synapse is located. The source of Ca^2+^ are L-type voltage-gated calcium channels in the model, Ca_v_1.2 and Ca_v_1.3, since L-type channels have been shown to mediate inhibitory plasticity in some neurons [66, 67]. In other neurons, multiple signals trigger inhibitory plasticity, such as R-type calcium channels important for LTP in layer 5 pyramidal neurons in the visual cortex, or cross-talk from Ca^2+^ influx through NMDA receptors from excitatory synapses [14, 68]. We did not include Ca^2+^ from R-type channels or NMDA receptors since both allow significant Ca^2+^ influx only at higher membrane voltages, and as such are not good indicators of synaptic activity at lower voltages. Also, they have specific roles in plasticity, based on which more detailed learning rules can be constructed in future studies.

Since [Ca^2+^]_i_ is a fast signal, we based the plasticity rule on a filtered variable of [Ca^2+^]_i_, called *F*, whose dynamics is slower and “smoother” than that of [Ca^2+^]_i_. It can be thought of as the concentration of a postsynaptic molecule whose activity level is dependent on [Ca^2+^]_i_ and which represents the average level of [Ca^2+^]_i_ in the time frame *τ*_*F*_.

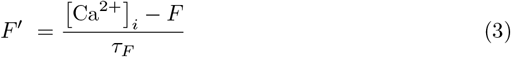

Using a filtered value of [Ca^2+^]_i_ makes the learning rule much less dependent on the fast variations in the calcium signal Fig. 1A_3_. However, in the previous version of the inhibitory plasticity rule used in [17], the maximal value of [Ca^2+^]_i_ was used, instead, showing that the rule is not critically dependent on filtering [Ca^2+^]_i_. Also, [Ca^2+^]_i_ from all voltage-gated calcium channels in the SPN model (not just L-type channels) was used in [17], suggesting that the results in this study will likely still hold if different sources of Ca^2+^ are used.

#### First type of simulations: control of neuronal firing rate

In the BCM rule, the function that updates the weights based on the indicator of synaptic activity, which in our case is the variable *F*, can be any nonlinear function that changes sign around the threshold level of activity [11]. To construct such a function, we have used a product of two sigmoidal functions, and the resulting inhibitory plasticity kernel used for controlling the neuronal firing rate is shown in Fig. 3A and given with Eqs. 4-6. The change in synaptic weight is given with Eq. 4:

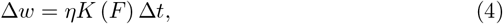

where *K* is the plasticity kernel, with its dependence on the indicator variable *F* written out explicitly, *η* is the learning rate, and Δ*t* is the simulation time step. The plasticity kernel is given with Eqs. 5 and 6:

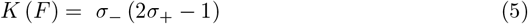

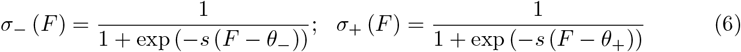

where *s* is the steepness of the sigmoidal curves (which is the same for the downward and upward slopes), *θ*_+_ is the plasticity threshold below which LTD is triggered, and above which LTP is triggered, and *θ*_−_− *c* is the minimal threshold level of *F* required for LTD to occur. *θ*_−_ is the half-maximum point of the downward-sloping part of the kernel, modeled with *σ*_−_ in Eq. 6, and *c* depends on the steepness parameter *s* in Eq. 6. The plasticity kernel becomes vanishingly small (close to 0) for values of *F* smaller than *θ*_−_− *c*, ensuring that a certain calcium activity is required for LTD. The kernel ranges from -1 to 1, and the learning rate *η* scales it so that changes in synaptic weight are done in small steps. (This *η* is different from the one in the NMDA receptor gating function in Eq. 2.) We set weights to 0 if a weight update causes them to become negative.

#### Second type of simulations: control of supralinear dendritic integration

In the second type of simulations, when the inhibitory rule is used for equalizing the dendritic response amplitudes, only homosynaptic plsaticity is used according to the same kernel as in the first type of simulations, given with Eqs. 4-6 and Fig. 3.

When the rule is used for gating or inhibiting supralinear dendritic responses, homosynaptic plasticity is formulated with an inverted kernel shown in and Fig. 7A and given with

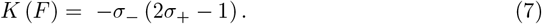

Also, for the inverted kernel, metaplasticity, or the so-called sliding threshold in the BCM rule, which modifies the value of *θ*_+_, is implemented according to the simple kernel in Fig. 7B and given with Eqs. 8 and 9. With this kernel *θ*_+_ follows the maximal value of *F* achieved during each pattern presentation. When *θ*_+_ reaches that value (usually after many update steps), synaptic plasticity stops. Because we use the maximal value of *F*, updates to the synaptic weights and the plasticity threshold are not done continuously (as in the first type of simulations), but during a 50 ms time window which starts 550 ms after a pattern is presented. During these 550 ms, the maximal value of *F* is reached.

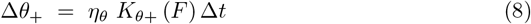

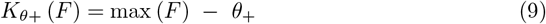

When heterosynaptic plasticity is added to work together with the inverted kernel for homosynaptic plasticity, it operates according to the same kernel as in Eqs. 4-6. The two kernels for homo- and heterosynaptic plasticity are shown together in Fig. 9A_1_. They have opposite shapes, resulting in opposite plasticity outcomes in the active and inactive inhibitory synapses. (Homosynaptic plasticity is triggered in the active synapses and heterosynaptic plasticity is triggered in the inactive ones). The usage of opposite shapes for the plasticity kernels is motivated by existing experimental results on heterosynaptic plasticity in inhibitory synapses [14, 19, 69, 70]. In these experiments, activating excitatory synapses while nearby inhibitory synapses are inactive produces opposite plasticity outcomes in both synapse types (eLTP and iLTD) [19, 70]. On the other hand, the same stimulation protocol used without activating any excitatory synapses produces eLTD and iLTP [19]. The differences in plasticity outcomes rely on differences in the evoked [Ca^2+^]_i_, which is higher when the stimulation protocol is paired with uncaging glutamate at an excitatory synapse, and lower when the stimulation protocol is used alone (without activating an excitatory synapse) [19]. This is consistent with different plasticity outcomes being triggered when [Ca^2+^]_i_ is below or above a threshold, as is captured by the BCM-type plasticity kernels that we use. However, experiments have shown molecular crosstalk between excitatory and inhibitory plasticity using various other signaling molecules apart from Ca^2+^ [1, 14]. We have not implemented such crosstalk between excitation and inhibition explicitly. Instead, the shape of the plasticity kernels determines the plasticity outcomes. Also, strictly speaking, to obtain the outcome described in [19], heterosynaptic plasticity in our case should instead be implemented with the kernel in Eq. 7, so we have not modeled the particular experimental findings in [19], but use them as motivation to formulate the oppositely-shaped plasticity kernels.

In the simulations with heterosynaptic plasticity, metaplasticity is also present, modifying the threshold *θ*_+_ in both kernels for homo- and heterosynaptic plasticity simultaneously, meaning that the two kernels use the same threshold *θ*_+_ (Figs. 9A_2_ and 9A_3_). We have tried two formulations of metaplasticity. The first is the same linear kernel as Eqs. 8 and 9, Fig. 3C (shown again in Fig. 9B_1_). However, this formulation causes inhibitory weights to grow without bounds, because of which we have used a second formulation with a nonlinear (quadratic) kernel, given with Eq. 10 and shown in Fig. 9B_2_. In fact, for stable synaptic weights, the BCM theory requires that *θ*_+_ is a nonlinear function of *F* [11].

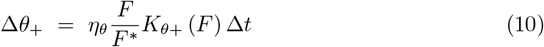

In Eq. 10, the amplitude of *F* is normalized by the maximal attainable value of *F* (denoted as *F*^*\**^) generated by an excitatory cluster of 20 synapses when no inhibition is present. This ensures that the quantity *F/F*^*\**^ is a value between 0 and 1, causing larger threshold updates only when the local Ca^2+^ activity is high.

To summarize, there are six parameter values in the inhibitory plasticity rule, shown in Table 2, followed by some notes on the chosen values. The parameter *s* determines the slope of the rising and falling parts of the kernel function. In the simulations for control of the neuron’s firing rate (the first type of simulations), where the weight updates are done continuously throughout the simulation, it is important that the slope is not too steep. If it is, the weight dynamics around the threshold *θ*_+_ exhibit oscillations because the weight updates are done in large steps, making *F* continuously over-/undershoot the threhsold *θ*_+_. The same holds for the learning rate *η* - large values can make large updates in synaptic weight, making *F* oscillate around *θ*_+_. This will translate to oscillations in the synaptic weights. In the simulations for control of supralinear dendritic integration (the second type of simulations), on the other hand, where weight updates are done for a brief period after the pattern presentation, this is not an issue, and both the slope and learning rates can be larger.

**Table 2.**
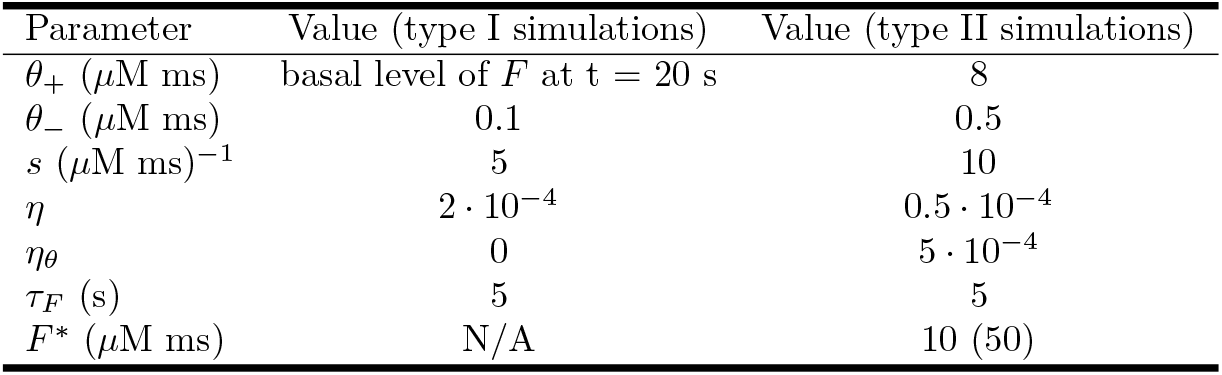
The parameters in the inhibitory plasticity rule. The value 50 for *F*^*\**^ is used in the simulations with the nonlinear feature binding problem.

#### Excitatory plasticity rule

In the last two figure of the Results section, in order to demonstrate that inhibitory plasticity can gate supralinear dendritic integration in a more dynamic setting, we have assumed that the clustered excitatory synapses are also plastic. We use a simple basal level of *F* at t = 20 s 0.1 plasticity rule based on the synaptic machinery in corticostriatal synapses onto direct-pathway SPNs (dSPNs) [21, 45]. Briefly, in the dSPNs, LTP occurs when dopamine stimulates the D_1_ dopamine receptors (D_1_Rs) and there is significant calcium influx from NMDA receptors [46, 71, 72]. LTD, on the other hand, occurs when no or little dopamine is bound to the D_1_Rs, and there is significant influx from L-type voltage-gated Ca^2+^ channels [61, 73, 74]. The plasticity rule for excitatory synapses is given with:

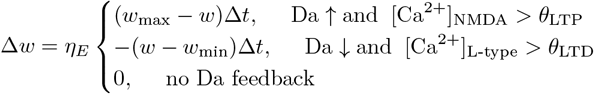

During LTP the weight can increase to a maximal weight of *w*_max_, and during LTD it can decrease to a minimal weight of *w*_min_, with a learning rate of *η*_*E*_. LTP happens after a dopamine peak is delivered and the NMDA Ca^2+^ concentration is higher than a threshold level *θ*_LTP_. LTD happens when a dopamine pause is delivered and the L-type Ca^2+^ concentration is higher than a threshold level *θ*_LTD_.

We have also incorporated a simple metaplasticity rule for the thresholds *θ*_LTP_ and *θ*_LTD_:

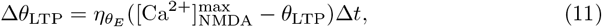

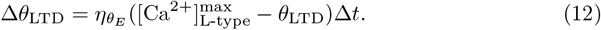

Both thresholds are updated when LTP or LTD occurs. With this metaplasticity rule, each of the thresholds follow the latest [Ca^2+^]_NMDA_ and [Ca^2+^]_L-type_ levels. Similarly to the inhibitory plasticity rule in the second type of simulations, the maximal [Ca^2+^]_NMDA_ and [Ca^2+^]_L-type_ levels attained during the presentation of each pattern are used. Briefly, metaplasticity in the LTP threshold prevents weakened synapses to participate in LTP easily, while metaplasticity in the LTD threshold protects strengthened synapses from easily undergoing LTD. This excitatory plasticity rule is demonstrated in detail in Figure 4 - figure supplement 2 of [21]. The parameters for the rule are given in Table 3.

**Table 3.**
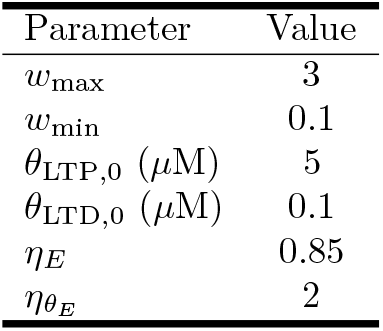
The parameters for the excitatory plasticity rule. *θ*_LTP,0_ and *θ*_LTD,0_ are the initial values of the thresholds in Eqs. 11 and 12.

Lastly, we briefly comment on the differences between this simple excitatory rule and the one developed in [17]. Both are based on the signals necessary for corticostriatal synaptic plasticity in dSPNs [45]. However, the aim with the excitatory rule in [17] was to study to what extent the NFBP can be solved with excitatory plasticity alone (while using a biologically-based learning rule). In this study, we focus only on the roles of inhibitory plasticity, so we use a simple excitatory plasticity rule that on its own cannot solve the NFBP. The two rules differ not only in complexity, they are also qualitatively different, despite being based on the same experimental results listed above. In the rule in [17], LTP happens within a region of [Ca^2+^]_NMDA_, and below and above this region no LTP occurs. Metaplasticity only affects the LTP process, and it does this by shifting the position of this region in an opposite direction of the evoked [Ca^2+^]_NMDA_. In this simple excitatory rule, both LTP and LTD happen above a respective threshold, and metaplasticity affects both thresholds so that they move towards the evoked [Ca^2+^]_NMDA_ and [Ca^2+^]_L-type_ levels (i.e. metaplasticity affects both the LTP and LTD process). Also, the rule in [17] has no explicitly defined limits on the synaptic weights as in this rule - it is meant to “find” synaptic weights that solve the NFBP.

#### Calcium dynamics

Intracellular calcium arising from NMDA receptors and the L-type voltage-gated channels Ca_v_1.2 and Ca_v_1.3 accumulates into two separate pools. Ca^2+^ extrusion is modeled with a calcium pump and with one-dimensional time decay. The calcium pump follows the implementation in [75]. The dynamics of the [Ca^2+^]_NMDA_ pool model were manually tuned to match the [Ca^2+^] amplitudes and durations reported in [76]. Spatial calcium diffusion was not implemented in this neuron model. As mentioned above, only Ca^2+^ from L-type channels is used in the inhibitory synaptic plasticity rule. The equation describing [Ca^2+^]_i_ is

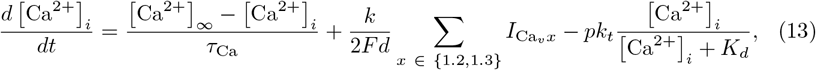

where *τ*_Ca_ is the [Ca^2+^]_i_ decay constant, *F* is the Faraday constant, *d* is the depth of the submembrane shell whel Ca^2+^ accumulates, *k*_*t*_ is the catalytic activity of the pump, *K*_*d*_ is the pump dissociation constant for Ca^2+^, and *k* and *p* are phenomenological parameters used in [75] to balance Ca^2+^ influx and efflux. The values for these parameters are given in Table 4.

**Table 4.**
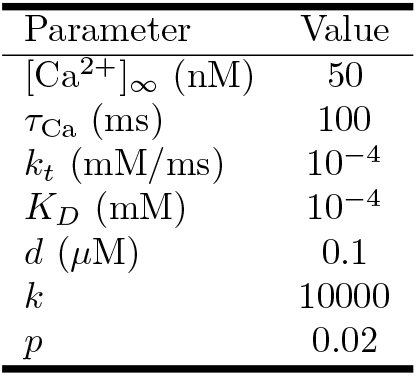
The parameters for the L-type Ca^2+^ pool model.

#### Glutamate spillover

In the last two figures of the Results section, where the clustered excitatory synapses are plastic, we have used the nonlinear feature binding problem as an example task for the neuron to solve using both excitatory and inhibitory plasticity. This task requires pronounced supralinear dendritic responses in order to be performed by the neuron. For this reason we used a glutamate spillover model to generate plateau potentials, detailed in [25]. According to this model, extrasynaptic NMDA receptors, located in the dendritic shaft which underlies the spines of the clustered excitatory synapses, are activated when the summed excitatory synaptic weights in the cluster are larger than a glutamate threshold level (here equal to 20 synapses, each with weight of at least 0.75). This evokes an all-or-none plateau potential which significantly depolarizes the soma and can drive somatic firing together with the background synaptic inputs. We also note that the glutamte threshold in this study is different from the values used in [25] and [17]. Without more studies on the conditions under which glutamate spillover occurs, the glutamate threshold remains a free parameter in the model.

#### Calcium diffusion

Also, in the last two figures of the Results section, where excitatory synapses are plastic, we have implemented calcium diffusion for [Ca^2+^]_NMDA_, the signal necessary for eLTP. We did this because without diffusion, the amplitude of [Ca^2+^]_NMDA_, which is used in the excitatory learning rule, does not show a monotonous increase for larger dendritic depolarizations. Instead, when the dendritic voltage approaches the NMDA receptor reversal potential (during a plateau, for example), the smaller NMDA current causes smaller [Ca^2+^]_NMDA_ amplitudes than those obtained for lower dendritic voltages (Fig. 9 in [21]). During learning, as excitatory synapses are strengthened, this effect causes the maximal [Ca^2+^]_NMDA_ amplitude to suddenly fall below the threshold for LTP, *θ*_LTP_, and prematurely stop learning in the excitatory synapses. To avoid this, we implemented axial calcium diffusion for [Ca^2+^]_NMDA_ according to the most detailed calcium diffusion model for SPNs in [76] (a description of the implementation is given in [21]). With axial calcium diffusion the amplitude of [Ca^2+^]_NMDA_ becomes proportional to the dendritic voltage that evokes it (Fig. 9C in [21]), and the excitatory synaptic weights reach at their maximal levels during learning (Figs. 11F and 11E, F).

We note that not implementing axial diffusion for [Ca^2+^]_NMDA_ does not affect the results in this study. This can be seen in S5 Fig which shows simulations for the symmetric input configurations in the NFBP performed with a pool model of [Ca^2+^]_NMDA_ according to Eq. 13, and using the parameters in Table 4. The excitatory weights do not reach their maximal values (S5 Fig, panels E and F), but the perfomance on the NFBP shows the same trend as in Fig. 11B where axial diffusion is implemented (S5 Fig, panel B, although less pattern presentations have been used to save computational resources).

#### Measuring performance on the nonlinear feature binding problem

We have quantified how well the neuron learns to solve the nonlinear feature binding problem with a performance score, and here we describe what the score means. As described below, there are four possible input patterns in the nonlinear feature binding problem. The neuron needs to learn to spike to only two of the patterns (which we call relevant patterns), and remain silent for the other two (irrelevant patterns). A score of 50% also means that the neuron is correct only half of the time. An example of such a score is when the SPN is silent for all four patterns, or spikes for all four patterns. Similarly, a score of 100% indicates it always spikes for the relevant patterns, and is always silent for the irrelevant patterns. A score of 75% is exemplified with the situation where the neuron is silent for the two irrelevant patterns, and spikes for only one of the relevant patterns. The threshold score for solving the task is 87.5%, where the SPN would also spike for the other relevant pattern at least half of the time.

## Acknowledgements

We acknowledge the use of Fenix Infrastructure resources, which are partially funded from the European Union’s Horizon 2020 research and innovation programme through the ICEI project under the grant agreement No. 800858. Simulations were also performed on resources provided by the National Academic Infrastructure for Supercomputing in Sweden (NAISS) at PDC KTH partially funded by the Swedish Research Council through grant agreement no. 2022-06725. This study was supported by the Swedish Research Council (VR-M-2020-01652), the Swedish e-Science Research Centre (SeRC), EU/Horizon 2020 No. 101147319 (EBRAINS 2.0 Project) and the European Union’s Research and Innovation Program Horizon Europe under grant agreement No. 101137289 (the Virtual Brain Twin Project).

## Supporting information

**S1 Fig. The effect of the inhibitory input frequency on the IO function**.

The input frequency of the inhibitory inputs is varied from 1.5 Hz to 3.0 Hz in steps of 0.1 Hz. The excitatory synaptic strength is 100 pS, and the inhibitory synaptic strength is 1.5 nS.

**S2 Fig. Homosynaptic inhibitory plasticity shows instability of the synaptic weights for long learning times**. (A) The input configuration is the same as in Fig. 6. (B) In each dendrite, all patterns are eventually inhibited. (C) The evolution of inhibitory synaptic weights (left column) and the thresholds *θ*_+_ (right column) during learning. Once max (*F*) falls below *θ*_+_ for the (two) strongest pattern(s) in each dendrite, a positive feedback loop is triggered where the inhibitory synapses for that (those) pattern(s) are increased, suppressing the evoked max (*F*) even further. This keeps the value of max (*F*) in the positive region of the kernel, persistently increasing inhibitory synapses. *θ*_+_ is shown with lines and max (*F*) with dots, only for the first synapse in each cluster (ovelapping dots are not highighted). (D) The average change in dendritic depolarization after learning. Positive values indicate increased dendritic responses, and negative values indicate reduced dendritic responses. All patterns are inhibited after learning. Results are averages of 128 trials, where in each trial the clusters are placed on three randomly chosen dendrites out of a set of 9 dendrites. Also, in each trial there are a total of 1800 pattern presentations arriving in random order. The difference is calculated between the mean of the last 10 and the first 10 appearances of a given pattern.

**S3 Fig. All possible input configurations that can solve the NFBP containing up to 3 features per dendrite**. Taking into account that no two dendrites in the SPN are precisely the same (i.e. are not symmetric), there are 18 such unique input configurations. In the learning simulations, in each input configuration the dendrites are additionally innervated with inhibitory clusters for all four features.

**S4 Fig. All possible input configurations containing 4 features in one or both dendrites**. Taking into account that no two dendrites in the SPN are precisely the same (i.e. are not symmetric), there are 13 such unique input configurations.

**S5 Fig. Targeted inhibition can solve the NFBP in difficult (symmetric) cases also when no axial calcium diffusion for [Ca**^**2+**^]_**NMDA**_ **is implemented**. In this figure, the pool model for [Ca^2+^]_NMDA_ given with Eq. 13 was used instead. (A) The input configuration with all four features present on two dendrites used in this figure. Inhibitory synapses target only certain features. (B) Performance on the NFBP averaged across all input configurations with four features on at least one dendrite, shown in S4 Fig. 19 trials are performed for each input configuration, for a total of 247 trials. In each trial, synapses are placed on two randomly chosen dendrites from a set of 9 dendrites. Not using axial diffusion for [Ca^2+^]_NMDA_ still allows for the NFBP to be solved when inhibition is used. Without inhibition, on average the same pattern is stored in both dendrites and the NFBP is not solved. (C) Voltage traces from the dendrites and soma for each pattern before and after learning, for the input configuration in (A) containing both excitatory and inhibitory synapses. (D) The same plot as in (C) when only excitatory synapses are present. The neuron also spikes for irrelevant patterns. (E) Evolution of excitatory and inhibitory synaptic weights in the two dendrites for the input configuration in (A). Excitatory synapses do not reach their maximal values. (F) Evolution of excitatory synaptic weights when no inhibitory synapses are present in the input configuration in (A). Excitatory synapses also do not reach their maximal values. In this figure *w*_max_ = 2, plateaus are generated with a lower glutamate threshold when 20 synapses reach weights of at least 0.5, and there are 5 spikes arriving per inhibitory synapse.

**Acknowledgments**

We acknowledge the use of Fenix Infrastructure resources, which are partially funded from the European Union’s Horizon 2020 research and innovation programme through the ICEI project under the grant agreement No. 800858. Simulations were also performed on resources provided by the National Academic Infrastructure for Supercomputing in Sweden (NAISS) at PDC KTH partially funded by the Swedish Research Council through grant agreement no. 2022-06725.

